# EpiAlign: an alignment-based bioinformatic tool for comparing chromatin state sequences

**DOI:** 10.1101/566299

**Authors:** Xinzhou Ge, Haowen Zhang, Lingjue Xie, Wei Vivian Li, Soo Bin Kwon, Jingyi Jessica Li

## Abstract

The availability of genome-wide epigenomic datasets enables in-depth studies of epigenetic modifications and their relationships with chromatin structures and gene expression. Various alignment tools have been developed to align nucleotide or protein sequences in order to identify structurally similar regions. However, there are currently no alignment methods specifically designed for comparing multi-track epigenomic signals and detecting common patterns that may explain functional or evolutionary similarities. We propose a new local alignment algorithm, EpiAlign, designed to compare chromatin state sequences learned from multi-track epigenomic signals and to identify locally aligned chromatin regions. EpiAlign is a dynamic programming algorithm that novelly incorporates varying lengths and frequencies of chromatin states. We demonstrate the effcacy of EpiAlign through extensive simulations and studies on the real data from the NIH Roadmap Epigenomics project. EpiAlign is able to extract recurrent chromatin state patterns along a single epigenome, and many of these patterns carry cell-type-specific characteristics. EpiAlign can also detect common chromatin state patterns across multiple epigenomes, and it will serve as a useful tool to group and distinguish epigenomic samples based on genome-wide or local chromatin state patterns.

## INTRODUCTION

All tissue and cell types, such as embryonic stem cells (ESCs), terminally differentiated tissues, and cultured cell lines, are maintained and controlled by epigenomic regulation and gene expression programs (1, 2, 3). An epigenome encodes information of chemical modifications to DNA and histone proteins of a genome, and such modifications may result in changes to chromatin structures and genome functions. Epigenomic information is represented by multi-track signals, including DNA methylation, covalent histone modifications, and DNA accessibility, all of which are measured genome-wide by high-throughput sequencing technologies such as Bisulfite-seq, ChIP-seq and DNase-seq (4). In recent years, multiple international consortia, including the Encyclopedia of DNA elements (ENCODE) (5), the NIH Roadmap Epigenomics Mapping Consortium (6, 7), and the International Human Epigenome Consortium (8), have generated large-scale high-throughput epigenome sequencing datasets for a broad spectrum of tissue and cell types, offering an unprecedented opportunity for studying multiple levels of epigenetic regulation across diverse cell states. Specifically, the NIH Roadmap project has released public epigenomic data of 127 human tissue and cell types (7). This database contains a total of 2,804 genome-wide epigenomic datasets, including 1,821 histone modification datasets, 360 DNase datasets, and 277 DNA methylation datasets.

A series of computational methods, including ChromHMM (9), Segway (10), GATE (11), TreeHMM (12), STAN (13), EpiCSeg (14), Spectacle (15), IDEAS (16), and GenoSTAN (17), have been developed to build a genome-wide chromatin state annotation, where distinct chromatin states have demonstrated diverse regulatory and transcriptional signals (18, 19, 20). In these methods, each epigenome is segmented into non-overlapping regions, and a single-track chromatin state sequence is constructed by compressing the multi-track epigenetic activities (e.g., DNA methylation and histone modifications) in various ways. For example, ChromHMM assigns discrete chromatin state labels to genomic regions based on signals of multiple epigenetic markers using a hidden Markov model (9). The predicted chromatin states have shown strong biological relevance and wide applicability in genomic research, e.g., the identification of enhancers and promoters (20). Given a chromatin state annotation constructed by any of these methods, genomic regions of the same chromatin state are expected to have both consistent epigenomic patterns and similar regulatory functions.

Based on existing chromatin state annotations, previous work has studied similarities and differences of human tissue and cell types in terms of epigenomic signals in specific functional genomic elements (e.g., promoters and enhancers), as well as the tissue and cell specificity of these elements, using the Pearson correlation coefficients (7, 21) or a newly developed epigenome overlap measure (EPOM) (22). The aforementioned methods have shed significant insights into our understanding of gene regulation on a global scale, i.e., how promoters and enhancers regulate target genes in diverse tissue and cell types. However, former epigenome comparative studies failed to effectively incorporate the sequential information of chromatin states, which, however, we believe are highly likely to contain critical information on gene regulatory mechanisms.

The comparison of DNA/RNA or protein sequences is based on the sequential information of nucleotides or amino acids. Many sequence alignment methods have been developed over the past decades to measure the similarity between sequences. Earlier work such as the Needleman-Wunsch algorithm (23) and the Smith-Waterman algorithm (24) use dynamic programming to search for the best global or local matches between two sequences. With the development of these algorithms, sequence alignment tools have become indispensable in almost all modern biological research. They are powerful not only in studies that focus on comparing sequences, such as evolutionary studies, but also in query-database retrieval studies, which aim to find regions from a large database that are similar to the query sequence of interest. However, there is no alignment algorithm designed to assess the epigenetic similarity of long genomic regions, such as gene regions and long non-coding regulatory regions. A main challenge lies in the multi-track nature of epigenomic signals. On the one hand, substantial information would be lost if we calculate a scalar value (e.g., the mean signal averaged over multiple 25 bp windows) to represent the signal of a long genomic region per track per tissue/cell. On the other hand, if we directly analyze the original data (a signal value per 25 bp window per track per tissue/cell), we would need to evaluate the similarity of large matrices to compare genomic regions. Specifically, the matrix of a region has the dimensions as the number of 25 bp windows in the region the number of tracks. Given that different regions almost certainly have different region lengths thus they have matrices of different dimensions, how to evaluate their similarity is a non-trivial task. In addition, we also need to consider the fact that a long region often contains multiple functional genomic elements with varying lengths. Hence, a reasonable approach is to compare two long regions based on their chromatin state patterns learned from multiple-track epigenomic signals. Motivated by the fact that chromatin state sequences provide a biologically meaningful one-track interpretation of multi-track epigenomic signals (9), we reduce the challenging question of comparing long multi-track epigenomic signals to a simpler task of comparing two chromatin state sequences.

Given the fast accumulation of large-scale epigenomic datasets generated in recent years, biological researchers are in great need of a new bioinformatic tool to efficiently retrieve genomic regions similar to an interested query region in terms of epigenomic signals. Motivated by the enormous successes of sequence alignment algorithms in comparing nucleotide and protein sequences (25), here we propose a novel computational method, Epigenome Alignment (EpiAlign), to compare two genomic regions by aligning their chromatin state sequences. To the best of our knowledge, EpiAlign is the first pairwise alignment-based method that investigates the sequential patterns of chromatin states and studies the epigenome similarity based on the patterns. EpiAlign compares two chromatin state sequences by calculating a local alignment score. It also allows the search of genomic regions (i.e., “hits”) whose chromatin state sequences are similar to those of a query region. Aligned chromatin state sequences are expected to have similar biological functions. EpiAlign is flexible in performing the chromatin state sequence alignment either within an epigenome, i.e., a tissue or cell, or between two epigenomes. From the alignment results of EpiAlign, users can identify common chromatin state patterns to investigate the functional relationship of interested genomic regions.

## METHODS

The EpiAlign algorithm aims to find an optimal local alignment between two chromatin state sequences. Our algorithm development is motivated by the classic Smith-Waterman Algorithm (24). We design the mismatch and deletion score functions based on the weight of each chromatin state in each sequence. We first apply a chromatin state annotation method (e.g. ChromHMM (9)) to encode multi-track epigenomic signals into single-track chromatin state sequences, whose different states are represented by different labels. Second, we compress consecutive occurrences of the same state into a state label. For example, a chromatin state sequence `abbcc` is represented by a compressed state sequence *S* = `abc`. EpiAlign then performs a local alignment between two genomic regions based on their compressed state sequences. The motivation of adding a compression step lies in the fact that most uncompressed (chromatin state) sequences contain long stretches of a single chromatin state, mostly the quiescent/low state (see Supplementary section 2), and including such length information would dominate the alignment result, a scenario that is often undesirable, because the purpose of alignment is to find similar chromatin state patterns composed of more than one state. The compression step allows EpiAlign to focus more on chromatin state patterns instead of a single chromatin state that spans a long genomic region. We use an example to demonstrate the effectiveness of adding the compression step to address this issue: in the brain sample E071, when we applied EpiAlign with the compression step, the brain-specific gene *NRG3* has the best alignment with another brain-specific gene *GRIA1*, among all the protein-coding genes. This result is reasonable as both genes are brain-specific and highly expressed in brain samples. However, as these two genes have vastly different lengths (*NRG3* is three times longer than *GRIA1*) and their chromatin state sequences have long stretches of the quiescent/low state, they are poorly aligned when we applied EpiAlign without the compression step. This result indicates that the compression step, which condenses the epigenetic information encoded in chromatin state sequences, is necessary and effective for finding similar and biologically meaningful chromatin state patterns. Additionally, aligning uncompressed sequences is much more time-consuming (20 times more computation time on average) than aligning their compressed counterparts. Therefore, adding the compression step also increases the computational efficiency of EpiAlign. In the following text, unless specified, all the chromatin state sequences refer to the compressed state sequences.

### Modified Smith-Waterman Algorithm for Chromatin State Sequence Alignment

Given two chromatin state sequences *S*_1_ and *S*_2_, we characterize a possible alignment between *S*_1_ and *S*_2_ through a set of triplets 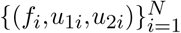, where *N* denotes the total number of aligned basepairs (including matches, mismatches, and gaps), *f*_*i*_ gives the alignment status between two chromatin states whose positions are *u*_1*i*_ and *u*_2*i*_ in *S*_1_ and *S*_2_, respectively. We may equivalently write this set of triplets as three equal-length sequences: *F* = *f*_1_*f*_2_ … *f*_*N*_, *U*_1_ = *u*_11_*u*_12_ *u*_1*N*_, and *U*_2_ = *u*_21_*u*_22_ … *u*_2*N*_. Specifically, *f*_*i*_ ∈ {m,n,d_1_,d_2_}denotes one of the four possible alignment status between two chromatin states: `m` for match, `n` for mismatch, d_1_ for deletion in *S*_1_, and d_2_ for deletion in *S*_2_. If *f*_*i*_ = `m`, there is a match between the *u*_1*i*_-th state of *S*_1_ and the *u*_2*i*_-th state of *S*_2_; if *f*_*i*_ = `n`, there is a mismatch between the *u*_1*i*_-th state of *S*_1_ and the *u*_2*i*_-th state of *S*_2_; if *f*_*i*_ = `d`_1_, the *u*_1*i*_-th state of *S*_1_ is aligned to nothing in *S*_2_ (*u*_2*i*_ is set to 0); if *f*_*i*_ = d_2_, the *u*_2*i*_-th state of *S*_2_ is aligned to nothing in *S*_1_ (*u*_1*i*_ is set to 0). In an example with *S*_1_ = `abca` and *S*_2_ = `aba`, if we consider an alignment 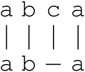, then *F* = mmd_1_ `m`, *U*_1_ = `1234`, and *U*_2_ =`1203`. Please note that the two chromatin state sequences *S*_1_ and *S*_2_ may have different lengths. Also given *S*_1_ and *S*_2_, it is possible to have more than one alignment results, i.e., sets of 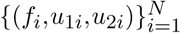.

Now we define the alignment score function *H*(*•*) as:

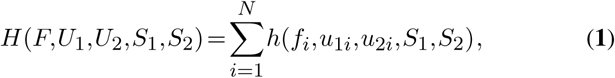

where *h*(*f*_*i*_,*u*_1*i*_,*u*_2*i*_,*S*_1_,*S*_2_) denotes the score of the alignment status *f*_*i*_ between the *u*_1*i*_-th state in *S*_1_ and the *u*_2*i*_-th state in *S*_2_. Specifically,

- *h*(`m`,*u*_1*i*_,*u*_2*i*_,*S*_1_,*S*_2_)=MF(*u*_1*i*_,*u*_2*i*_,*S*_1_,*S*_2_);
- *h*(`n`,*u*_1*i*_,*u*_2*i*_,*S*_1_,*S*_2_)=MF(*u*_1*i*_,*u*_2*i*_,*S*_1_,*S*_2_);
- *h*(`d`_1_,*u*_1*i*_,*u*_2*i*_,*S*_1_,*S*_2_)=DF(*u*_1*i*_,*S*_1_);
- *h*(`d`_2_,*u*_1*i*_,*u*_2*i*_,*S*_1_,*S*_2_)=DF(*u*_2*i*_,*S*_2_);

We will formally define the matching function MF (.) the mismatching function NF(*•*) and the deletion function and the deletion function DF(·) later in this section. To summarize, the function *h*(·) takes a form that depends on the value of its first argument *f*_*i*_.

Then we consider the alignment problem as an optimization problem where the goal is to find the optimal alignment 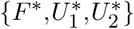 that maximizes the alignment score *H*:

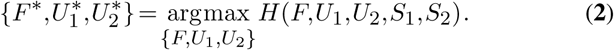

This optimization problem can be approached by dynamic programming, an algorithm that iteratively maintains and updates a matrix *M* that stores dynamic alignment results. The matrix element *M*_*k,l*_ is the maximal alignment score of the two subsequences 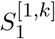 and 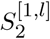, where 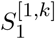 denotes the first *k* states of *S*_1_ and 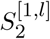 denotes the first *l* states of *S*_2_. Let *n*_1_ and *n*_2_ be the length of *S*_1_ and *S*_2_, respectively. We update the matrix *M* using the following rule.

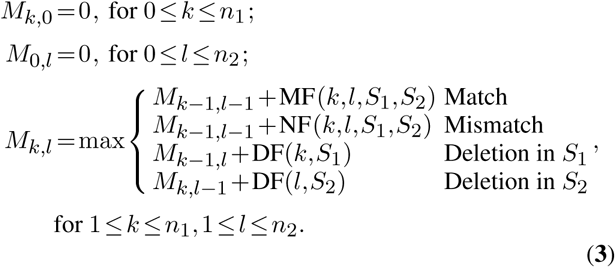

The algorithm described in Equation (**3**) achieves the global alignment, but we instead consider the local alignment approach in practice since the local alignment would prefer long continuous alignments with small proportion of mismatches, which are more likely to contain the common patterns of interest. In contrast, global alignment would prefer patterns containing overly scattered short alignments separated by gaps. To achieve the goal of local alignment, we propose the following approach to modify the dynamic programming algorithm.

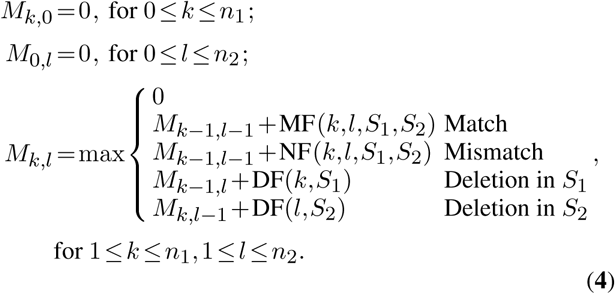

The alignment score of EpiAlign is 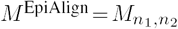.

### Chromatin State Weights

To define the specific forms of the matching function MF(·),the mismatching function NF (•),and the deletion function DF(·), we first introduce a weight function *W* (*k,S*), which describes the weight of the *k*-th state in sequence *S*. The weights can be used to distinguish chromatin states of different importance if we have prior knowledge that some states have more significant biological functions than others at certain positions. We design two sets of weights: (1) **equal weights** mean that all states are treated equally with the same weight 1 in sequence *S*, i.e., *W* (*k,S*)=1,*k* = 1,…, |*S*|; (2) **frequency-based weights** assign larger weights to less common chromatin states (see Supplementary section 1 for details), motivated by the fact that some uncommon states are likely strong indicators of biological functions.

With the weights defined above, we specify the matching function, the mismatching function, and the deletion function as:

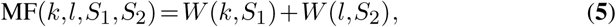

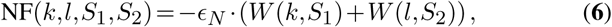

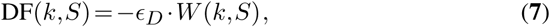

where ϵ_*N*_ and ϵ_*D*_ are the penalty parameters for a mismatch and a deletion in the alignment, respectively. In EpiAlign, ϵ_*N*_ and ϵ_*D*_ can be tuned by users, and the default values are 1.5 and 1, respectively. The choice of ϵ_*N*_ and ϵ_*D*_ values depends on how “local” users would like the result to be, i.e., if we set a larger ϵ_*N*_ or ϵ_*D*_ value, it means that we penalize more on a mismatch or a gap in the alignment, and the final best alignment result will be shorter or more local. Figure 1 shows the workflow of EpiAlign.

**Figure 1.**
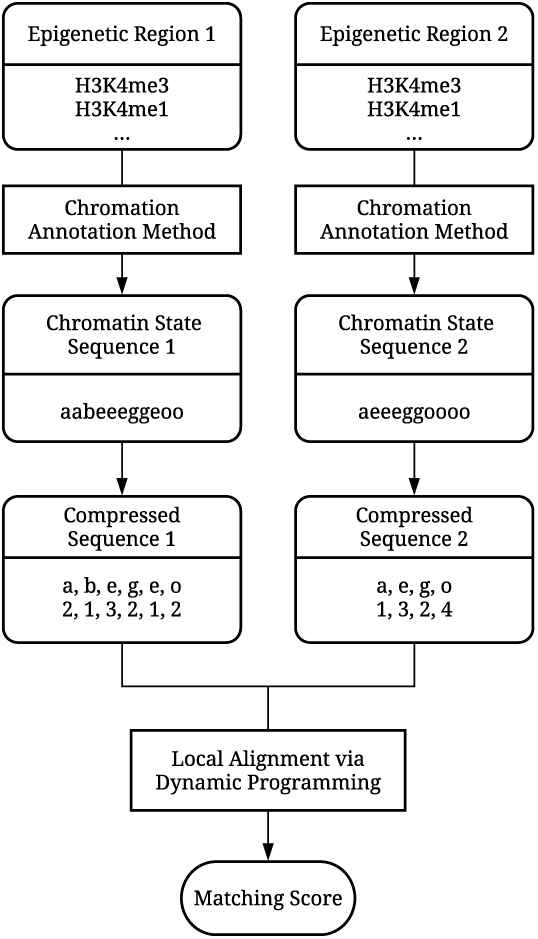
Workflow of the EpiAlign algorithm.

## RESULTS

We demonstrate in three aspects that EpiAlign is a useful tool for investigating sequential patterns of chromatin states. First, we demonstrate that EpiAlign can identify common chromatin state patterns within the same epigenome or across different epigenomes. Second, we investigate biological interpretation of the common chromatin state patterns found by EpiAlign. Third, as a technical verification, we show that EpiAlign is able to distinguish real epigenomes from randomized epigenomes. We also demonstrate the superiority of EpiAlign over a naïve method that compares two chromatin sequences only based on chromatin state frequencies. We conduct the above analysis using simulation and real case studies based on the Roadmap epigenomic database (7). In this paper, we use the chromatin state sequences annotated by ChromHMM, which has been well recognized to provide an informative compression of multi-track epigenomic signals into a chromatin state sequence (7, 9, 22). It is worth noting that our method is generally applicable to chromatin state sequences annotated by other methods.

In this paper, for most analysis, we selected ESC, heart and brain samples from the Roadmap dataset as representative examples. The reason is that among all the Roadmap tissue types, these three types are relatively better understood and have well-annotated tissue-specific genes(26).

### Vertical alignment: Comparison of Chromatin State Sequences of Protein-coding Genes across Epigenomes

EpiAlign is a powerful local alignment algorithm to quantify the similarity of two chromatin state sequences in terms of their aligned subsequences. Here we apply EpiAlign to compare chromatin state sequences of the same genomic region in different epigenomes, a strategy we define as the **vertical alignment**. The diversity of the same region’s chromatin state sequences represents epigenetic characteristics of various tissues and cell types. As epigenetic characteristics are known to have a strong association with gene expression characteristics (27), we expect that a cell-type specific gene, i.e., a gene specifically highly expressed in a cell type (26), should have similar chromatin state sequences in epigenomes of that cell type. In contrast, lower similarity is expected between two chromatin state sequences, one of that cell type and the other of another cell type (Supplementary Figures 3 and 4).

**Figure 2.**
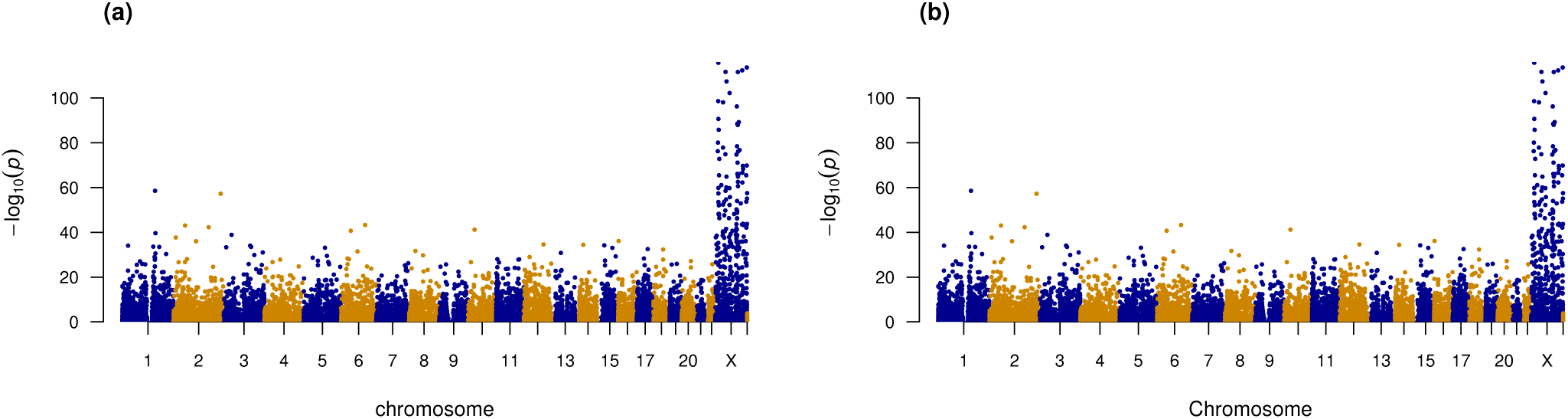
Alignment scores of chromatin state sequences of protein-coding genes within a sex vs. between sexes. We perform the two-sample one-sided Wilcoxon test between within-sex alignment scores and between-sex scores to quantify their differences: (a) Manhattan plot of *p*-values of the test between male-vs-male and male-vs-female alignment scores for all the protein-coding genes. (b) Manhattan plot of *p*-values of the test between female-vs-female and male-vs-female alignment scores for all the protein-coding genes. In the two comparisons, within-sex and between-sex alignment scores differ most significantly for genes on the X chromosome.

**Figure 3.**
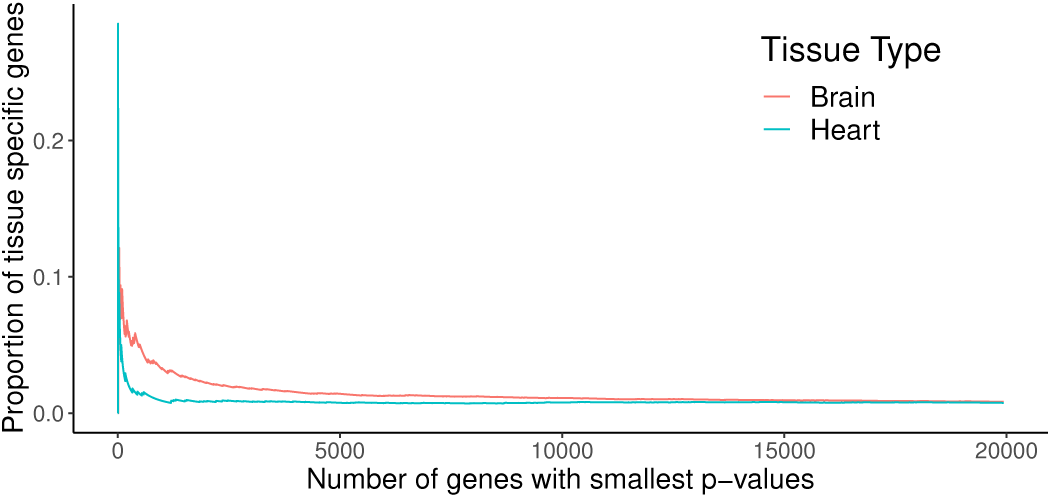
Brain and heart specific genes are enriched in the top differential genes that have significantly higher within-tissue alignment scores than between-tissue scores. The horizontal axis shows the number of top differential genes, and the vertical axis shows the proportion of tissue specific genes among the top differential genes.

**Figure 4.**
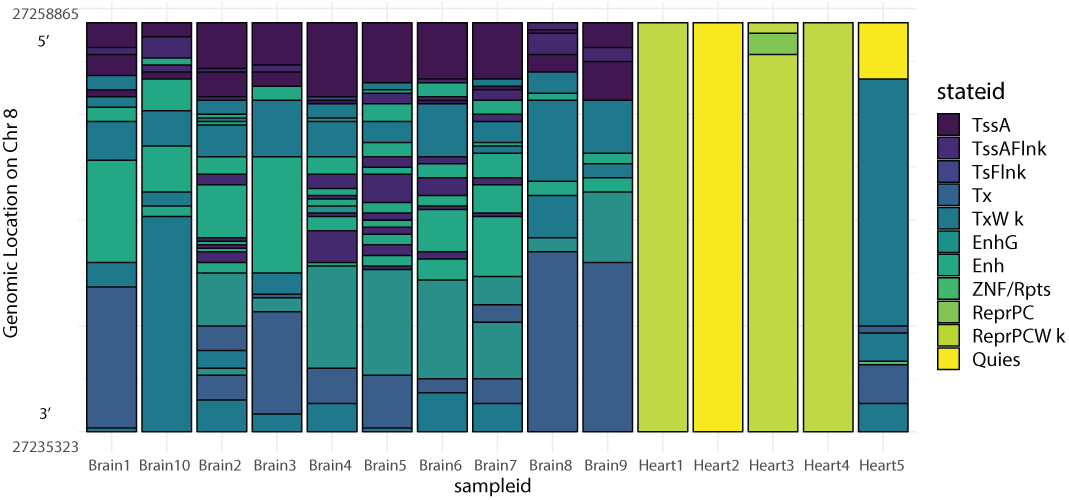
Chromatin state sequences of gene *STMN4* in all the 10 brain samples and the 5 heart samples. Different chromatin states are represented by different colors. The y-axis indicates the genomic locations of various chromatin states across these 15 samples.

In the first study, we divide the Roadmap epigenomes into two categories: 51 male samples and 38 female samples. In the second study, we compare the Roadmap epigenomes of two cell types: 10 brain samples and 5 heart samples. In both studies, we compare the chromatin state sequences for each of the 19,935 protein-coding genes between every pair of samples. (Note that we use all protein-coding genes in GENCODE v10 (28) that are compatible with the Roadmap database, with the exception of genes on chromosome Y.)

We obtain three sets of alignment scores: pairwise scores within male samples, pairwise scores between male and female samples, and pairwise scores within female samples. Since most genes on the X chromosome are associated with sex-linked traits, we expect to observe higher alignment scores between samples of the same sex than those between samples of different sexes. To quantify the difference between alignment scores, we perform the two-sample one-sided Wilcoxon test between male-vs-male scores and male-vs-female scores for each protein-coding gene. Studying the resulting p-values, we find that out of the top 200 genes that have the smallest p-values, 188 are X chromosome genes. (Figure 2(a)). This result suggests that the majority of the genes that exhibit greater within-sex similarity are sex linked, a reasonable finding that matches our expectation. The comparison between female-vs-female and male-vs-female alignment scores leads to a similar result (Figure 2(b)). These results together confirm that EpiAlign successfully distinguishes same-sex chromatin state sequences from different-sex ones, suggesting that EpiAlign outputs a reasonable similarity measure of chromatin state sequences.

We also investigate the 12 genes that are not on X chromosome among the top 200 genes with the smallest p-values (Supplementary Table 1). These genes are potentially sex linked. For example, *MFF* that controls mitochondrial fission has been reported to have to have sex-specific regulation (29). This result suggests that EpiAlign can serve as a useful tool for discovering genomic regions with certain epigenetic regulation of interest.

**Table 1.** Alignment scores of chromatin state sequences of protein-coding genes within a tissue (heart or brain) vs. between heart and brain. Displayed are the enriched GO terms in the top 200 significant genes identified by the Wilcoxon test between brain-vs-brain alignment scores and brain-vs-heart alignment scores. The top enriched GO terms are highly relevant to heart processes or brain processes (*: terms related with heart; **: terms related with brain).

| GO term | Description | P-value |
| --- | --- | --- |
| GO:0051891 | *positive regulation of cardioblast differentiation | 9.34E-8 |
| GO:0051890 | *regulation of cardioblast differentiation | 6.42E-7 |
| GO:0007416 | **synapse assembly | 5.82E-6 |
| GO:0003207 | *cardiac chamber formation | 5.83E-6 |
| GO:0060413 | *atrial septum morphogenesis | 1.72E-5 |
| GO:0006928 | movement of cell or subcellular component | 2.15E-5 |
| GO:0007409 | **axonogenesis | 2.98E-5 |
| GO:0071625 | vocalization behavior | 3.07E-5 |
| GO:0032990 | cell part morphogenesis | 4.63E-5 |
| GO:2000738 | positive regulation of stem cell differentiation | 6.36E-5 |
| GO:0060043 | *regulation of cardiac muscle cell proliferation | 6.99E-5 |
| GO:0097104 | **postsynaptic membrane assembly | 8.69E-5 |
| GO:0048812 | **neuron projection morphogenesis | 8.79E-5 |
| GO:0051705 | multi-organism behavior | 9.73E-5 |

In the second study, we investigate if EpiAlign can help identify cell-type specific genes, which were previously discovered from gene expression profiles (26), using only chromatin state sequences. We perform the two-sample one-sided Wilcoxon test between brain-vs-brain alignment scores and brain-vs-heart alignment scores for all the 19,935 protein-coding genes. We next perform the Gene Ontology (GO) enrichment analysis (30) on the top 200 genes that receive the smallest p-values in the Wilcoxon test (Supplementary Table 2). Here we choose the top 200 genes instead of setting a threshold on multiple-testing-adjusted p-values, because we found that the most commonly used threshold 0.05 led to a large number of significant genes. For our purpose of verifying that the top differentially aligned genes are biologically meaningful, choosing a smaller number of top ranked genes is a more reasonable approach. The top enriched GO terms (p-value < 0.0001) are highly relevant to heart/cardiac processes and brain processes (Table 1). Previously discovered 150 heart-specific genes and 166 brain-specific genes (26) are enriched in the top differential genes found by the Wilcoxon test, which have significantly higher within-tissue alignment scores than between-tissue scores. For example, 9 brain-specific genes and 4 heart-specific genes are in the top 100 differential genes (p-values < 10^−30^ in a hyper-geometric test). Figure 3 shows that top differential genes contain a higher proportion of tissue-specific genes. The above results indicate that EpiAlign is able to distinguish cell-type specific genes by assigning them higher alignment scores when comparing the epigenomes of their associated cell types. This again suggests that EpiAlign effectively captures chromatin state patterns in epigenomes.

**Table 2.**
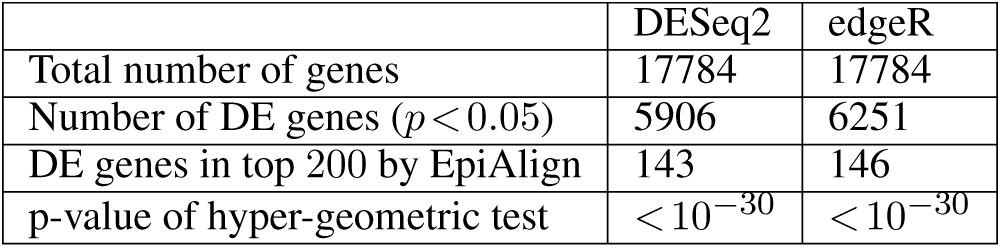
Comparison of the 200 genes with differential chromatin state sequences identified by EpiAlign and the differentially expressed (DE) genes identified by DESeq2 or EdgeR. DESeq2 and edgeR identify 5906 and 6251 DE genes between all 3 brain samples and all 4 heart samples from the 17,784 protein-coding genes in the Roadmap RNA-seq datasets. A hypergeometric test is used to check the significance of the enrichment of the top 200 genes identified by EpiAlign in the two sets of DE genes. The two resulting *p*-values are both significant.

|  | DESeq2 | edgeR |
| --- | --- | --- |
| Total number of genes | 17784 | 17784 |
| Number of DE genes ($p < 0.05$) | 5906 | 6251 |
| DE genes in top 200 by EpiAlign | 143 | 146 |
| p-value of hyper-geometric test | $< 10^{-30}$ | $< 10^{-30}$ |

To better illustrate how EpiAlign helps identify common chromatin state patterns, we study a brain-specific gene *STMN4*, which has the lowest p-value from our two-sample one-sided Wilcoxon test described above (brain-brain alignment scores vs. brain-heart alignment scores). Using it an example, we investigate the chromatin state sequences of *STMN4* in all brain and heart samples. From Figure 4, we observe that the brain samples share similar chromatin sequences; yet the common pattern in these sequences drastically differs from the chromatin state sequences in the heart samples. The fact that EpiAlign captured *STMN4* as the top differentially aligned gene shows that EpiAlign can successfully identify regions where chromatin state patterns diverge or conserve between cell types.

We also analyze the expression profiles of protein-coding genes. We use DESeq2 (31) and EdgeR (32) to do differential expression (DE) analysis between heart samples and brain samples on all the 17,784 protein-coding genes included in the Roadmap RNA-seq datasets. The results show a high consistency between the resulting differentially expressed genes and the differential chromatin state sequences found by EpiAlign (Table 2). This results further validate that the tissue-specific regions found by EpiAlign are biologically meaningful and reflect gene expression dynamics, and that EpiAlign will be a useful tool for identifying tissue-specific epigenomic regions.

### Horizontal Alignment: Analysis of Frequent Chromatin State Sequence Patterns within an Epigenome

Motivated by the fact that similar chromatin state sequences may encode similar biological functions, here we use EpiAlign to analyze frequent chromatin state sequence patterns within an epigenome. We introduce the “**horizontal alignment**,” which takes the chromatin state sequence of a region as the query and searches for its best hit except itself within an epigenome. We first divide a given epigenome into regions of 500 kb length, and then we align the chromatin state sequence of each region (i.e., the “query”) to those of other regions to find the best match. It is worth noting that the alignment scores of multiple query chromatin state sequences are not directly comparable. To normalize the alignment scores, we align every query chromatin state sequence to randomized chromatin state sequences, which serve as a negative control (see Supplementary section 3 for details). Then for every region, we define the normalized alignment score of its best hit except itself (when the region is used as the query) as its **horizontal alignment score**. A high score indicates that the region shares a highly similar and non-random chromatin state sequence with another region in the same epigenome, implying that the region’s chromatin state sequence pattern is likely biologically meaningful.

With horizontal alignment scores, we can represent every epigenome by a vector, whose length is the number of regions and whose entries are the regions’ horizontal alignment scores. As mentioned above, horizontal alignment scores measure whether their corresponding regions contain biologically meaningful chromatin state patterns, which are expected to be largely consistent across epigenomes of the same tissue. We use the Roadmap samples to calculate the horizontal alignment scores for all regions in all epigenomes. Then we represent every epigenome by a horizontal alignment score vector. To verify the biological meaning of the vector representation, we calculate the pairwise Pearson correlations between epigenomes and perform an average-linkage hierarchical clustering of epigenomes based on the (1–Pearson correlation) distance metric. The clustering result matches our expectation: samples from the same tissue are clustered together, confirming that the horizontal alignment scores are indeed consistent across the samples from the same tissue (Figure 5).

**Figure 5.**
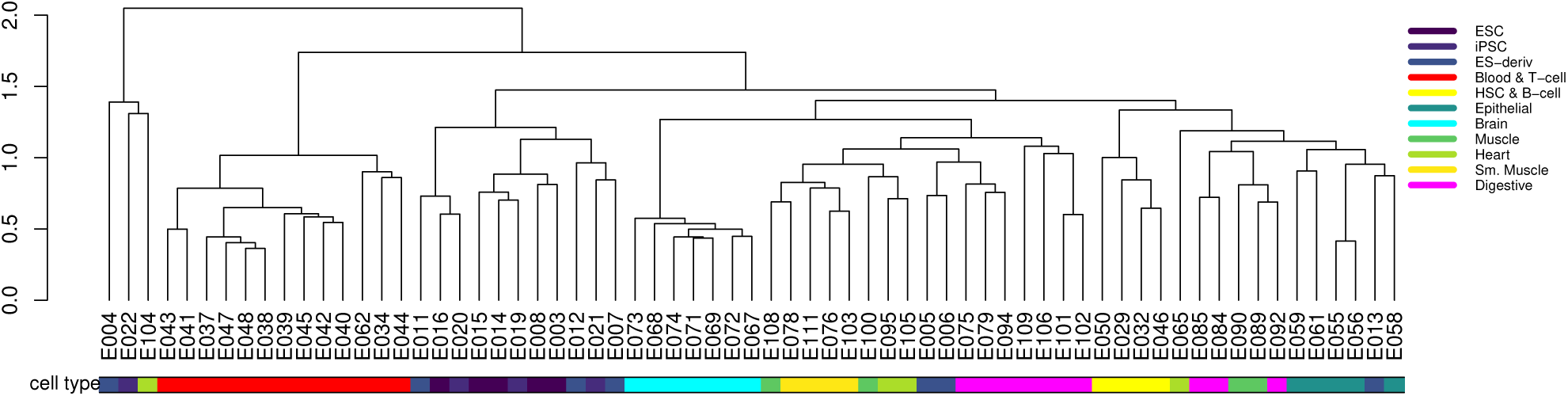
Clustering based on the correlation matrix of horizontal alignment scores of Roadmap epigenomes. Samples from the same tissue or cell type are clustered together, indicating that horizontal alignment scores are highly correlated between samples from the same tissue or cell type.

**Figure 6.**
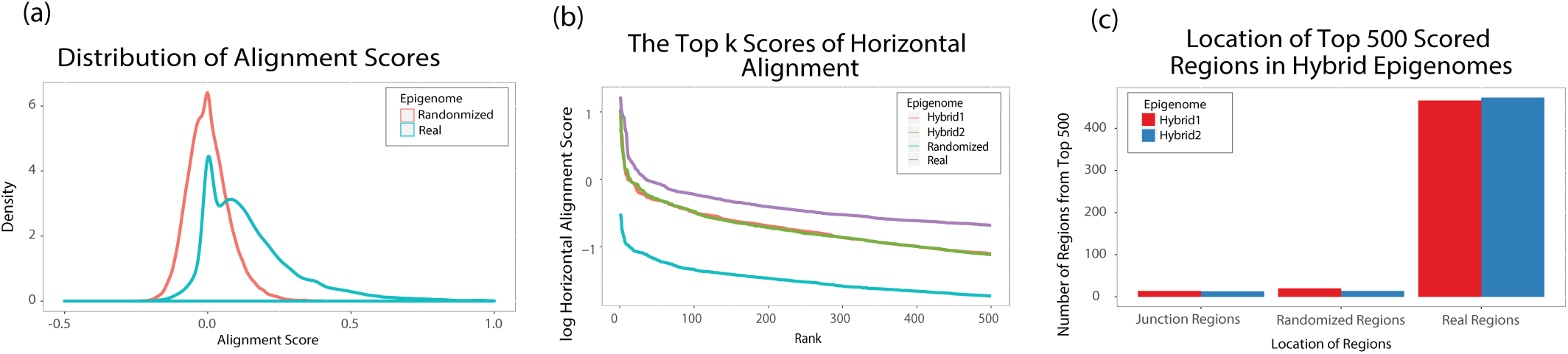
Horizontal alignment results of embryonic stem cell sample E003. (a) The distribution of horizontal alignment scores of regions in real and randomized epigenomes. (b)The top 500 highest horizontal alignment scores (log_10_ transformed) in real, randomized and hybrid epigenomes. Scores in the real epigenome are always the highest given the same rank. (c) Locations of the regions with the top 500 horizontal alignment scores in the two hybrid epigenomes. The three panels together indicate that the real epigenome contains non-random chromatin state sequential patterns captured by EpiAlign.

#### EpiAlign distinguishs real epigenomes from randomized ones

We further perform a simulation study to technically validate the efficacy of EpiAlign in terms of horizontal alignment. Our goal is to check if EpiAlign is able to distinguish real epigenomes from randomized epigenomes, which serve as a negative control. We calculate horizontal alignment scores using EpiAlign on all the 127 Roadmap samples based on the 15-state ChromHMM annotation. In addition to each real epigenome, we also generate a randomized epigenome and two hybrid epigenomes for comparison. Here the randomized epigenome is generated in the same way as in the normalization step for calculating horizontal alignment scores (see Supplementary section 3 for details). To contrast real and randomized epigenomes, we also generate a hybrid epigenome as a semi-negative control by mixing the real and randomized epigenomes of every chromosome, so that a hybrid epigenome is composed of alternating real regions and randomized regions. (see Supplementary section 4 for details)

We use an ESC (embryonic stem cell) sample (Roadmap ID E003) as an example and calculate horizontal alignment scores in four epigenomes: the real ESC epigenome, a randomized epigenome, and two hybrid epigenomes. We summarize the distributions of horizontal alignment scores in the real and randomized epigenomes in Figure 2(a). As expected, the regions in the real epigenome have an average alignment score higher than 0, while the average score of regions in the randomized epigenome is close to 0. For each of these four epigenomes, we find the top 500 non-overlapping regions with the highest horizontal alignment scores. As expected, the top regions in the real epigenome have scores significantly higher than those in the randomized and hybrid epigenomes (Figure 2(b)), an observation consistent with the fact that a high score indicates a region likely to have a biologically meaningful chromatin state pattern. Moreover, for hybrid epigenomes, almost all the top 500 regions are those generated from the real epigenome (Figure 2(c)), again confirming that real chromatin state patterns are more biologically meaningful than randomized patterns. Overall, our results suggest that EpiAlign can powerfully distinguish real biological epigenomes from randomized epigenomes.

#### Comparison of EpiAlign with alternatives

We further validate our EpiAlign algorithm with equal weights by comparing it with two alternative approaches. The first is a variant of EpiAlign using frequency-based weights, which are determined by the frequencies of chromatin states (see Supplementary section 1 for details). The second is a naïve alignment method, in which we first calculate the proportion of each chromatin state in two regions (chromatin state sequences) to obtain two proportion vectors *P*_1_ = (*p*_11_,*p*_12_,*…,p*_1*Q*_)^T^ and *P*_2_ =(*p*_21_,*p*_22_,*…,p*_2*Q*_)^T^, where *Q* is the number of unique chromatin states in the annotation(e.g., *Q* = 15 in this case). The naï ve alignment score is a similarity measure defined as 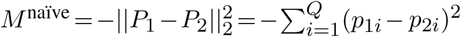. The naïve method directly compares two chromatin state sequences based on their state proportions, and it does not use a dynamic programming approach as does in EpiAlign. However, given that similar chromatin state sequences share similar frequency vectors, the naïve method is also a biologically meaningful approach.

Note that EpiAlign (with equal weights), the frequency-based variant of EpiAlign, and the naïve method do not have horizontal alignment scores on the same scale and cannot be compared directly, so we compare the three approaches by evaluating the biological meaning of the regions they find with high scores. Since gene regions are expected to share some common chromatin state patterns (i.e., promoter, transcription start site, transcribed region, and transcription ending site), a good alignment method is expected to assign high horizontal alignment scores to gene regions. In other words, genes expressed in a tissue are expected to have high horizontal alignment scores in the tissue’s epigenome. Hence, we design two evaluation criteria: one is the enrichment of known tissue-associated genes, i.e., the non-house-keeping genes highly expressed in a tissue (33), in regions with high alignment scores; the other criterion is the enrichment of annotated genes. The greater the enrichment, the better the alignment method. We apply each of the three approaches to do horizontal alignment and check the overlap between tissue-associated genes or annotated genes and each approach’s top-aligned regions, which receive the highest horizontal alignment scores. We perform this evaluation on 16 samples: 5 ESC, 4 heart and 7 brain samples. For each sample, we collect the top 500 regions with the highest alignment scores found by each approach and count the numbers of tissue-associated genes from Yang et al. (33) and annotated genes from Kent et al. (34) that overlap with these regions. From the results shown in Figure 7, we see that EpiAlign outperforms the naïve method in detecting annotated genes and tissue-associated genes. In addition, we observe that the frequency-based weights do not have apparent advantages over the equal weights, suggesting that we may use EpiAlign with equal weights as the default.

**Figure 7.**
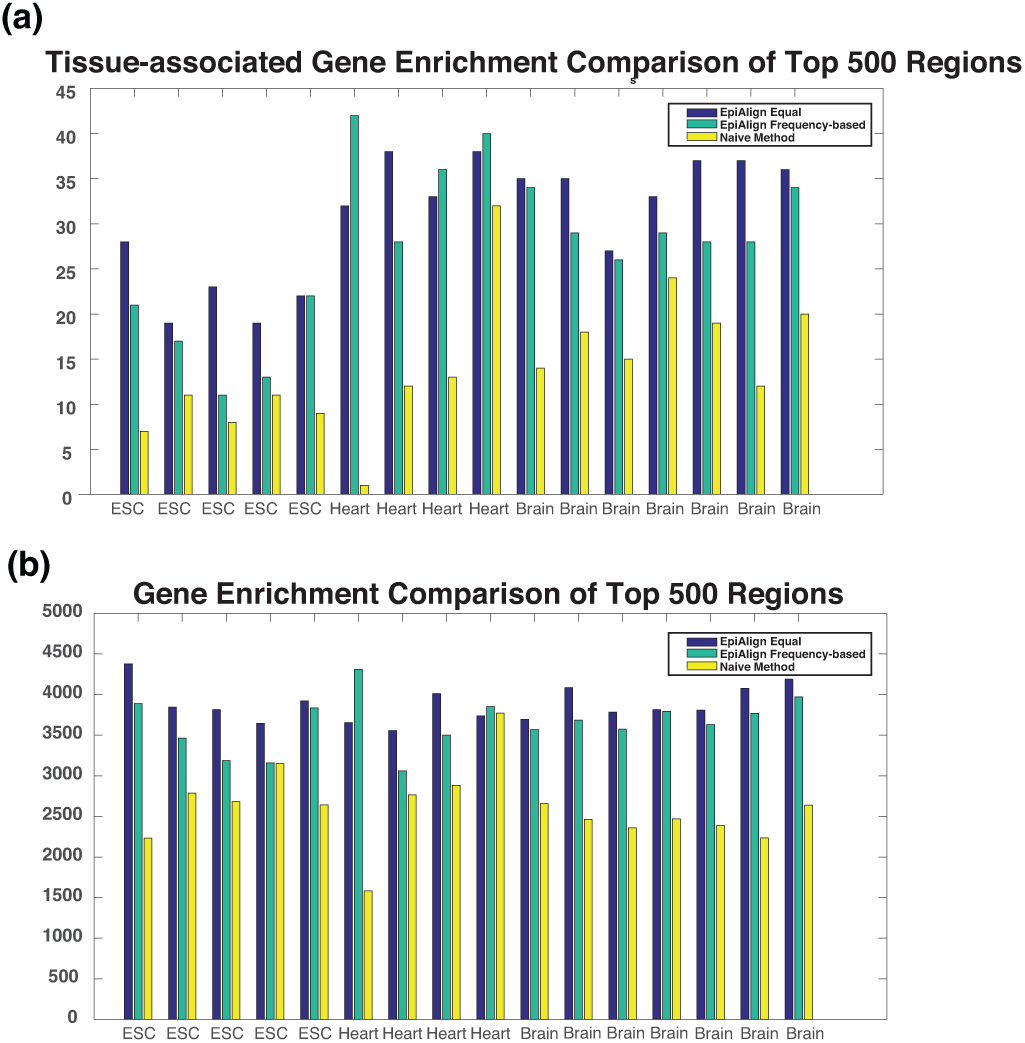
Comparison of EpiAlign, EpiAlign with frequency-based weights, and the naïve method using 16 Roadmap samples (5 ESC, 4 heart, and 7 brain samples from the 92 samples with 18-state ChromHMM annotation). (a) The number of tissue-associated genes that overlap with the top 500 regions with the highest horizontal alignment scores found by each approach. (b) The number of annotated genes that overlap with the same three sets of top 500 regions.

#### Motif Analysis

As a further investigation, we check if the regions with top horizontal alignment scores share any chromatin state patterns in common. We apply EpiAlign to perform horizontal alignment within the epigenome of the embryonic stem cell sample E003, and we select the top 200 regions with the highest horizontal alignment scores. To investigate whether common chromatin state patterns exist among these regions, we calculate the pairwise alignment scores between each pair of these top 200 regions. We normalize the pairwise alignment scores and store them in a 200 × 200 symmetric matrix **A**, whose (*i,j*)-th entry *A*_*ij*_ represents the normalized alignment score of regions *i* and *j* and is defined as

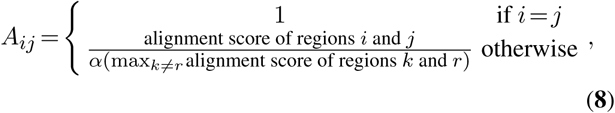

where *α*= 1.1 ensures that 0 *<A*_*ij*_ < 1 for all *i ≠ j*. We then define a distance matrix **D**, whose (*i,j*)-th entry is *D*_*ij*_ = 1 *A*_*ij*_. We then perform hierarchical clustering with average linkage on the top 200 regions based on **D**, and we display the clustering result in Figure 8.

**Figure 8.**
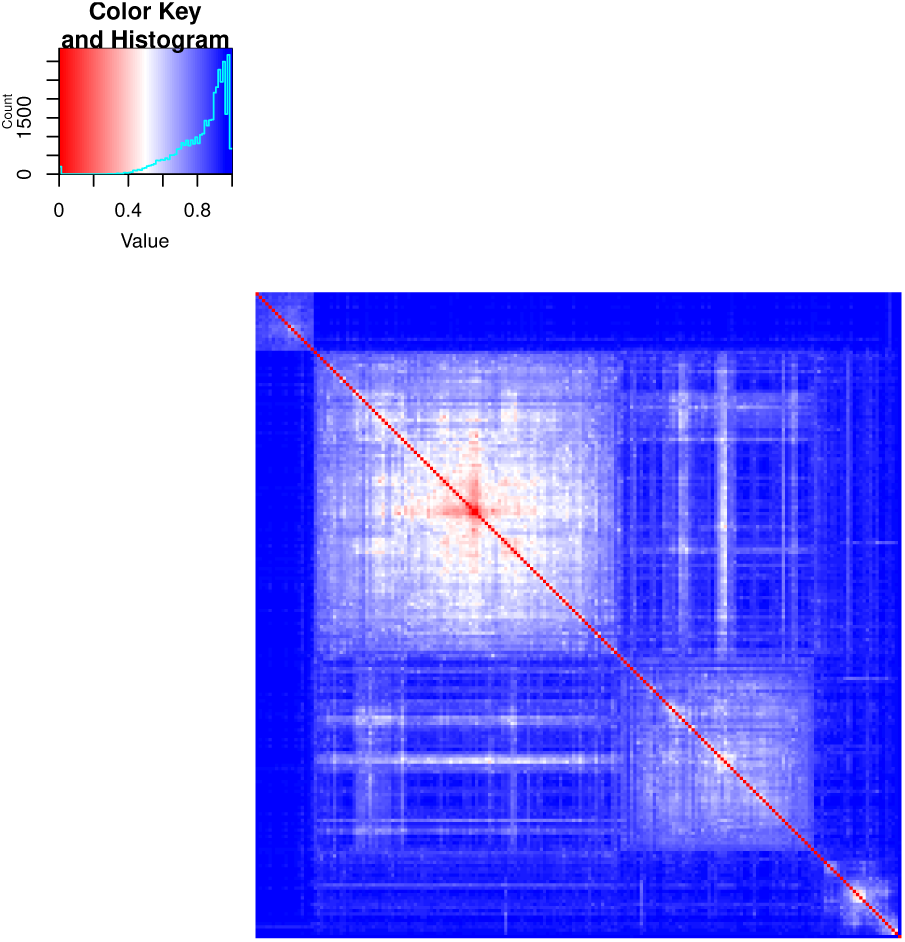
Heatmap of pairwise distances of the top 200 regions, identified by the horizontal alignment on embryonic stem cell sample E003. Based on the distance matrix **D**, the top 200 regions are grouped into 4 clusters by average-linkage hierarchical clustering.

From the heatmap in Figure 8, we see that the top 200 regions are well partitioned into four clusters, indicating that regions in the same cluster share similar chromatin state patterns. (Supplementary Table 3) We inspect each of these four clusters to identify its representative chromatin state patterns, which we refer to as *motifs* in the following text. For notation simplicity, we use alphabets “a” to “o” to denote chromatin states 1 to 15.

Using the motif-discovery tool MEME (35), we find that all the four clusters are characterized by certain motifs. As annotated by the 15-state ChromHMM model (36), the state “o” denotes the quiescent state and lacks a good biological interpretation, so we only consider the motifs without “o”. We find that cluster 1 is characterized by the “ihih”-repeat motif; cluster 2 is characterized by the “egeg”-repeat motif; cluster 3 is characterized by “eded” motif; cluster 4 is characterized by the “egeg” motif and “mlml” motif. Based on the ChromHMM annotation, the state “i” represents heterochromatin, while “h” represents ZNF genes and repeats. Since existing evidence shows that human heterochromatin proteins form large domains containing KRAB-ZNF genes (37), the “ihih”-repeat motif may represent functional non-coding regions. Since “d” denotes strong transcription, “e” denotes weak transcription, and “g” denotes enhancer, the “egeg”-repeat motif may be an evidence of transcriptional enhancers (38) and the “eded”-repeat motif may denotes transcriptional regions. In the “mlml”-repeat motif, “m” and “l” represent repressed polycomb and bivalent enhancer, respectively. Since polycomb-repressed genes have permissive enhancers that initiate reprogramming (39), the “mlml”-repeat motif may be an indicator of polycomb-repressed gene regions. All these results show that the motifs discovered from the frequent chromatin state patterns are biologically meaningful and EpiAlign can help identify common chromatin state patterns in epigenomes of specific biological conditions.

#### Cross-species application of EpiAlign

We further investigate the application of EpiAlign to comparing human and mouse chromtain state sequences. We use the epigenetic data from Yue et al. (2014), where mouse and human samples were used together to train a 7-state ChromHMM model(40). We investigate two liver samples, one from human and one from mouse. As homologous genes are expected to exhibit more similar functions than non-homologous genes(41), we expect to observe larger alignment scores between chromatin state sequences of homologous genes than those of non-homologous genes of similar sequence lengths. Our analysis is as follows. We first obtain mouse-human homologous gene pairs from Ensembl BioMart (Release 95) (42). We sort the mouse genes with lengths 200-400 kb by gene lengths and divide the homologous gene pairs into 12 groups each with 50 pairs, so that the mouse genes within a group have similar lengths. Within each group, we apply EpiAlign to each mouse-human homolog pair and each non-homolog pair. The results show that among the 12 groups, on average 16% the human genes have the highest chromatin state sequence alignment scores with their corresponding mouse homologs, suggesting that homologous genes tend to share similar epigenetic patterns. We also look at the GO terms of the homolog pairs that have the highest alignment scores in each group. The result (see Supplementary Table S4) shows that homologous genes with high alignment scores are also very similar in molecule functions and biological processes. The result also indicates that EpiAlign can identify homologous genes whose epigenetic patterns are more conserved in evolution, shedding new insights into translating scientific discoveries in mice into humans.

## WEBSITE

We have implemented the EpiAlign algorithm in an open-access software package, which is available at GitHub: https://github.com/zzz3639/EpiAlign

We have also created a user-friendly website to demonstrate the functionality of EpiAlign and visualize the alignment results of the Roadmap epigenomes: http://shiny.stat.ucla.edu:3838/EpiAlign.

The website includes two main features: cell-type alignment scores and pairwise alignment scores. For the cell-type alignment feature, users can browse the alignment score matrix for a given gene. The columns and rows of this symmetric matrix correspond to the 16 cell types, and each matrix entry is the average pairwise alignment score between the gene’s chromatin state sequences of the two corresponding cell types. For the pairwise alignment feature, users can select two gene regions and calculate the alignment score between their corresponding chromatin state sequences. Both features will help users investigate for a specific gene the similarity of its chromatin state patterns between Roadmap epigenomes or users’ custom epigenomic samples.

## DISCUSSION

In this article, we propose the EpiAlign algorithm for alignment of chromatin state sequences learned from multitrack epigenomic signals. We demonstrate that EpiAlign can be a powerful tool for studying the epigenetic dynamics along the same epigenome or across multiple epigenomes, based on both simulation and real data studies.

First, our current alignment results are based on ChromHMM, which learns and characterizes from multi-track epigenomic signals. There are also other tools for pattern discovery in chromatin structures, such as Segway (10), which constructs a dynamic Bayesian network instead of HMM, EpiCSeg (14), which uses natural numbers instead of binarized signals as used by ChromHMM, and IDEAS (16), which jointly characterizes epigenetic dynamics across multiple human cell types. It would be interesting to compare these tools with ChromHMM to analyze how the chromatin state annotation affects the alignment results of EpiAlign. If the output results of ChromHMM or other segmentation tools can be filtered or improved based on additional biological experiments, this can also help EpiAlign obtain more accurate and robust results. Besides, we find likely noisy ChromHMM annotations that need further biological validation (see Supplementary section 12). To account for such possible inaccuracy in chromatin state sequences, we may improve EpiAlign by incorporating the posterior probabilities of chromatin states output by ChromHMM into the calculation of alignment scores. Moreover, ChromHMM is an unsupervised algorithm that requires a pre-specified number of states; thus, its chromatin state labels may not be fully biologically meaningful. For example, some genomic regions would be assigned to different chromatin states given different numbers of states. This leads to additional noise in ChromHMM annotations. To account for such noise, we may correct chromatin state labels by using the sequential information in neighboring states.

Second, in the EpiAlign algorithm, an important step before alignment is the compression of the chromatin state sequences. Chromatin states of different regulatory functions can vary greatly in their lengths (43), but the length information itself is not always informative of the change of epigenetic marks along the genome. Specifically, the quiescent/low state often appear in extremely long stretches, whose lengths are not useful for comparing chromatin state sequences (see Supplementary Figure S1). Therefore, we add a compression step to capture and extract the dynamics of chromatin states among biological samples. We have also tested the pre-compression alignment algorithm, but it is not able to distinguish the randomized chromosome from the real one, suggesting that compression is necessary for detecting biologically meaningful chromatin state patterns. However, we realize that this compression step still has room for improvement. For example, several previous studies have shown that broad/sharp H3K4me3 domains have distinct functions (44, 45, 46), implying that the length information of certain chromatin states is important for vertical alignment that compares a region across samples. Future refinement of the compression step, or refinement of length information usage after compression, should consider multiple aspects: a chromatin state’s confidence (whether it is likely noisy) and importance (whether its length information is informative), as well as the analysis needs (vertical or horizontal alignment), among others.

Third, EpiAlign is essentially an unsupervised algorithm, but the flexibility of the weight function allows EpiAlign to incorporate prior knowledge into the alignment procedure by assigning different weights to different chromatin states. For example, the frequency-based weights lead the algorithm to favor the alignment of less frequent patterns compared to background patterns, which frequently exist along the epigenome. In practical applications, one may adjust the weight function to reflect the important elements in specific problems. For instance, the weight can incorporate the transcription start sites (TSSs) in genome annotation when transcriptional regulation is of particular importance.

Fourth, EpiAlign depends on two tuning parameters: ϵ_*N*_ and ϵ_*D*_ for penalizing mismatches and gaps in the alignment. Similar parameters are also necessary for classical alignment algorithms designed for DNA and protein sequences such as BLAST. For example, the ϵ_*D*_ in EpiAlign is analogous to the Gap Extend Penalty in BLAST. The NCBI BLAST, an online tool that implements the BLAST algorithm, sets the Gap Extend Penalty to 1 by default. In EpiAlign, we also set ϵ_*D*_ to 1 by default. In BLAST, a substitution matrix is used to score matches/mismatches, and multiple substitution matrices have been constructed for users to select based on alignment purposes. In EpiAlign, we set ϵ_*N*_ to 1.5, which is equivalent to a substitution matrix with diagonal entries as 1 and off-diagonal entries as −1.5. Given that the alignment of epigenetic sequences is new to this field, how to construct more specialized substitution matrices for chromatin states is an important future research question.

Finally, in some computationally efficient sequence alignment algorithms, hash tables or tree-based data structures are utilized to index the database, and these techniques have greatly increased the efficiency of query retrieval. EpiAlign can also benefit from similar techniques and further improve its computation efficiency.

Two other computational methods, EpiCompare (47) and ChromDiff (48), have been developed to compare chromatin states between samples. They test for the difference of a single chromatin state’s frequency in a genomic region between two groups of samples. EpiCompare restricts the region of interest to a 200 bp window, which corresponds to a single chromatin state output by ChromHMM. A useful functionality of EpiCompare is that it searches for the 200 bp windows where the specified chromatin state is enriched only under one condition. Compared with EpiCompare, ChromDiff is more flexible and allows the region to have any length greater than 200 bp. Another advantage of ChromDiff is that it normalizes the chromatin state frequencies to reduce the effects of confounding covariates. A common limitation of ChromDiff and EpiCompare is that they can only compare chromatin state frequencies between two conditions in the same genomic region, and they require multiple samples under each condition. In contrast, EpiAlign can perform pairwise alignment between any two chromatin sequences, either coming from the same genomic region in two samples or two different genomic regions in one sample. In other words, EpiAlign does not pose any restrictions on the choice of genomic regions or the sample size. Furthermore, EpiAlign has two unique advantages. First, it simultaneously uses the sequential information encoded in multiple chromatin states. Second, it outputs an alignment score that integrates this sequential information. Hence, EpiAlign enables horizontal alignment and query search, allowing us to extract chromatin state patterns that carry tissue-associated characteristics. These patterns are shown to be biologically meaningful in our motif analysis and have a strong capability in grouping epigenomic samples of the same cell type in horizontal alignment.

In terms of biological applications, the biggest strength of EpiAlign is its ability to identify common chromatin state patterns and how they are conserved or divergent between cell types. This strength will pave the way for identifying regulatory domains defined by combinatorial effects of strings of cis-elements. Specifically, the vertical analysis based on EpiAlign will reveal tissue-specific genes and regulatory regions that share common chromatin state patterns within a tissue type, and such patterns will serve as the basis of defining new regulatory domains. We have also demonstrated that EpiAlign has found meaningful chromatin state motifs. Besides, EpiAlign is able to distinguish tissue-associated genes. These results suggest the potential of EpiAlign as a useful bioinformatic tool to discover tissue-specific gene regulation. Moreover, the alignment scores calculated by EpiAlign can serve as a covariate when constructing functional genomic networks, thus allowing the network to incorporate similarities of chromatin structures as a factor. Further, EpiAlign applies to 3D genomic analysis to address the question if there are chromatin state patterns in regions with a specific 3D structure such as a loop.

## ACKNOWLEDGEMENTS

We deeply appreciate the insightful feedbacks from Prof. Jason Ernst at UCLA and Dr. Yucheng T. Yang at Yale University. We also thank Prof. Changshui Zhang at Tsinghua University for supporting H.Z.’s research.

