## Supplementary materials for "EpiAlign: an alignment-based bioinformatic tool for comparing chromatin state sequences"

#### 1 Frequency-based weights

The frequency-based weight of the $k$-th state in chromatin state sequence $S$ is $W^{Fb,w}(k,S)$, which is defined as:

$$W^{Fb,w}(k,S) = GS^w(k,S) \cdot RS^w(k,S) \cdot LS(k,S)$$

where $w$ is a window size parameter, $GS^w(k,S)$ is the two-state frequency score, $RS^w(k,S)$ is the one-state frequency score, and $LS(k,S)$ is the length score.

In the subsequence $S^{[k-w,k+w]}$, if neither of pattern $S^{[k-1,k]}$ or $S^{[k,k+1]}$ occur elsewhere in the subsequence, $GS^w(k,S) = 1$; if both patterns occur somewhere in the subsequence, $GS^w(k,S) = 4$; otherwise $GS^w(k,S) = 2$.

The one-state frequency score $RS^w(k,S)$ represents the frequency of each state and gives more frequent states smaller scores. For $S^{[k-w,k+w]}$, we rank the chromatin states in this window by their frequencies, from the highest to the lowest. Then

$$RS^w(k,S) = 1 + \frac{\text{rank}(S^{[k]}) - 1}{\text{number of unique states in } S^{[k-w,k+w]} - 1} \in [1,2].$$

In the compression process of EpiAlign, we compress consecutive occurrences of the same state into a state label. We also obtain an occurrence number for each state. For example, a chromatin state sequence `abbcc` is represented by a compressed state sequence $S =$ `abc` and a state occurrence sequence $L =$ `122`. The length score $LS(k,S)$ is based on the occurance sequence $L$. If $L^{[k]} = 1$, $LS(k,S) = 0.5$; otherwise $LS(k,S) = 1$.

#### 2 Average length of stretches of the same chromatin state

The motivation for doing compression before alignment comes from the fact that most uncompressed sequences contain long stretches of the same chromatin state. Here we use letter "a" to "o" to denote chromatin state 1 to 15 from the Roadmap 15-state annotation. We calculate the average length of consecutive same chromatin state for each state. From the results in Figure S1, we can see that the chromatin state "o", which means quiescent/low state, is much longer than other states before compression. As the length information of such a state is hardly biologically meaningful. The compression step is needed for addressing this issue by turning the focus onto the more biologically meaningful chromatin state patterns.

#### 3 Horizontal alignment scores

It is worth noting that the alignment scores of multiple query chromatin state sequences are not directly comparable. To normalize the alignment scores, we align every query chromatin state

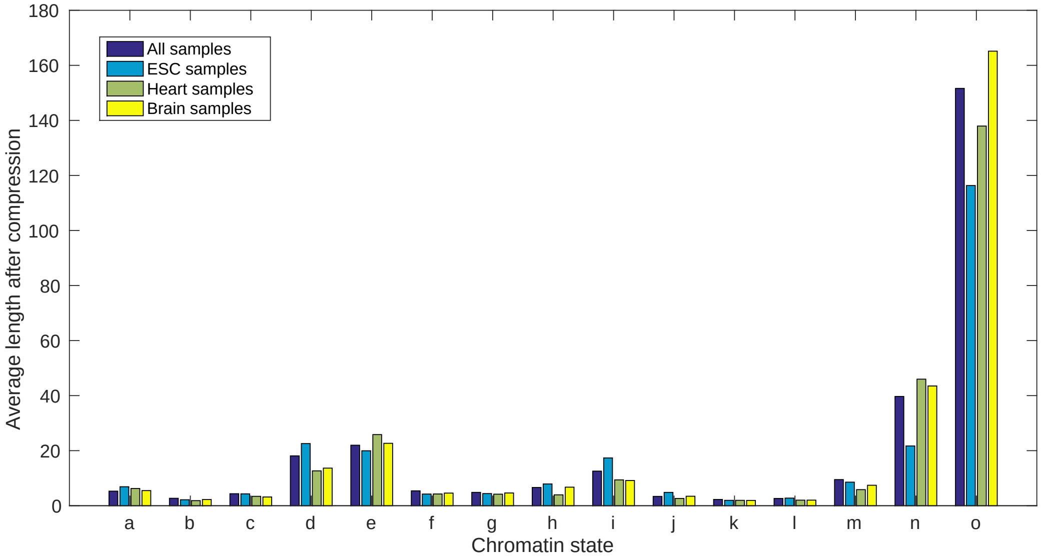


Figure S1: Average lengths of stretches of a single chromatin state in ESC, heart and brain samples. Letters 'a' to 'o' refer to chromatin states 1 to 15 in the Roadmap 15-state chromHMM annotation. For each state, we calculated the average length of its stretches, i.e., consecutive occurrences. It is obvious that the chromatin state 'o', i.e., the quiescent/low state, has much longer stretches than other states do.

sequence to randomized chromatin state sequences, which serve as a negative control. For each region in the real epigenome, this region is used as the "query" and aligned to each of the three randomized epigenomes to obtain its hit in that randomized epigenome. Here the randomized epigenomes have the same lengths as their real counterparts and are generated by the Markov rule, with a state transition probability matrix per chromosome based on the real epigenome. The alignment scores of the three hits are then averaged as the baseline score of this query region. We use $Q_i$ to denote the alignment score of region $i$'s hit in the real epigenome, $P_i$ to denote the baseline score of region $i$, and define the horizontal alignment score of region $i$ as $\frac{Q_i - Pi}{Pi}$. A high score indicates that the region shares a highly similar and non-random chromatin state sequence with another region in the same epigenome, implying that the region's chromatin state sequence pattern is likely biologically meaningful.

### 4 Generation of hybrid epigenomes

For every chromosome, we first divide both its real chromatin state sequence ("epigenome") and their randomized counterparts into non-overlapping regions of 50 million bp length. Then for the $i$-th region in the hybrid epigenome, its chromatin state sequence is set as the sequence of the $i$-th region in the real epigenome when $i$ is even, or as the sequence of the $i$-th region in the randomized epigenome when $i$ is odd. We can easily generate another hybrid epigenome if we switch the odd and even regions.

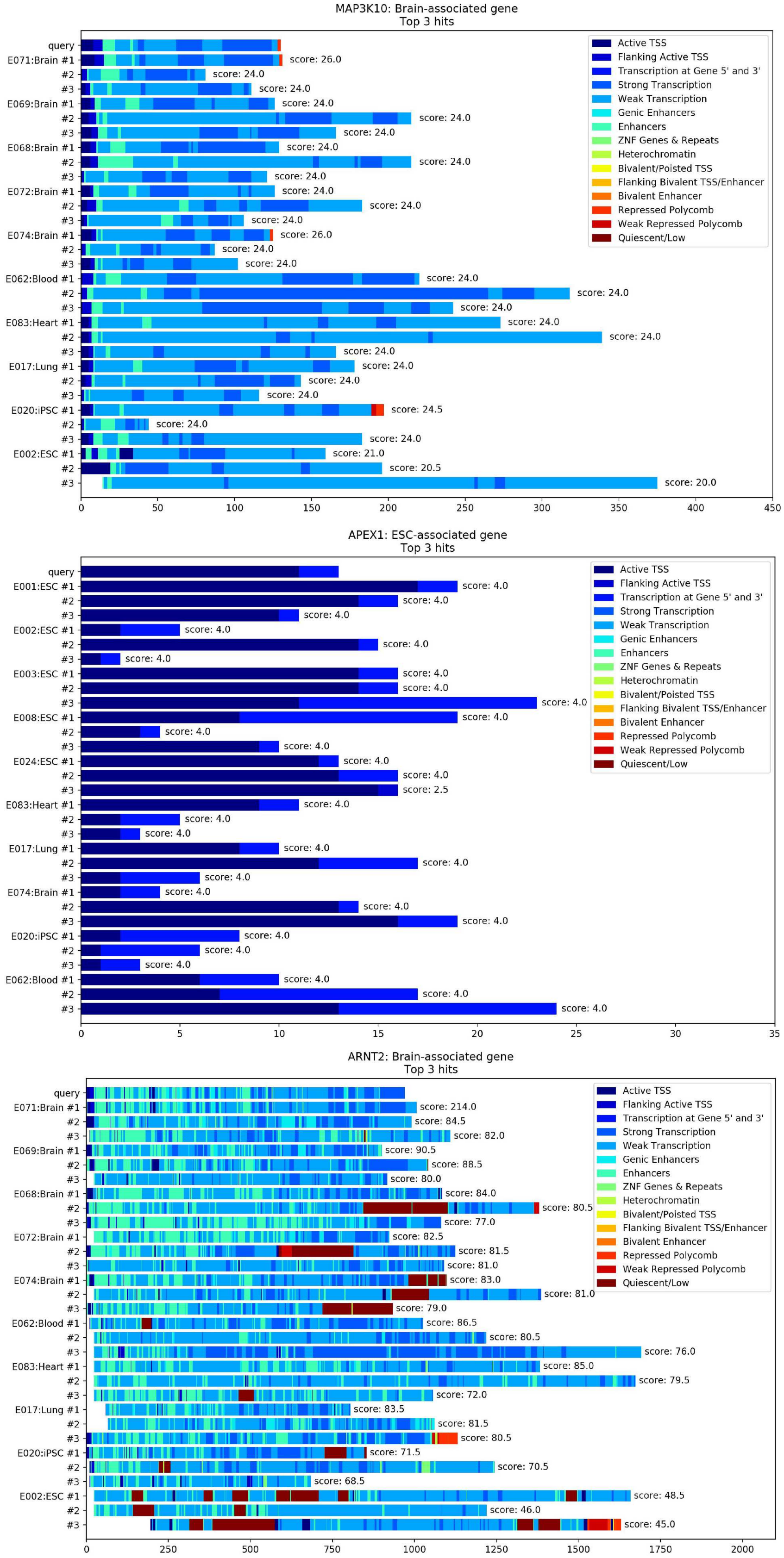


Figure S2: Top hits of 3 different queries in epigenomes from various tissues. The queries are chromatin state sequences of a brain-associated gene *MAP3K10*, an ESC-associated gene *APEX1*, and a brain-associated gene *ARNT2*.

### 5 Visualization of top hits in horizontal alignment

Figure S2 shows that when we use the chromatin state sequences of gene regions as queries, the best hits (regions that have the highest horizontal alignment scores with the query) reported by EpiAlign are very similar to the query in terms of chromatin state patterns.

### 6 Comparison of uncompressed sequences and compressed sequences in Vertical Alignment

We also use the vertical alignment to justify our choice of aligning compressed chromatin state sequences instead of original uncompressed sequences. We repeated the vertical alignment analysis on all brain-specific genes and all heart-specific genes among the brain and heart samples, using the uncompressed sequences instead of the compressed chromatin state sequences. Then we performed the same two-sample one-sided Wilcoxon test between brain-vs-brain alignment scores and brain-vs-heart alignment scores on these selected genes, and we denote the resulting p-values as uncompressed p-values. Next we compare these uncompressed p-values with their corresponding p-values we obtain previously based on the compressed sequences. We counted the number of significantly different genes, which have a p-value less than 0.05 after Bonferroni correction, from both analyses. From compressed sequences, 112 out of 327 tissue-specific genes are significant and from uncompressed 120 out of 327 are significant. We also conducted two-sample Wilcoxon test between the original p-values and p-values from uncompressed sequences. The result shows no significant difference in distribution (p-value = 0.509). These results show that the resulting p-values from the compressed sequences are very similar to those from the uncompressed sequences. Considering that alignment of uncompressed sequences is much more time-consuming (takes 20 times more time than compressed sequences), the compression step makes the alignment algorithm more effective.

### 7 Examples of vertical alignment on tissue-associated genes

Since epigenetic marks carry important regulatory information relevant to cell differentiation, chromatin states learned from these marks should also contain cell-type characteristic patterns. For a tissue-associated gene, we should expect to observe significantly higher similarity of chromatin states within its associated cell type than the counterpart similarity between cells of other cell types. We implement vertical alignment on the Roadmap dataset on some tissue-associated genes [1]. Taken an ESC-associated gene *ANAPC1* as an example, we use the alignment scores calculated by EpiAlign to compare the similarity of *ANAPC1*'s chromatin state sequences in different cell types. We first extract the chromatin state sequence of *ANAPC1*'s chromatin region from in each epigenome. Then, we use EpiAlign to calculate the alignment scores of these chromatin state sequences between each pair of the 127 epigenomes, resulting in $8,001$ pairwise alignment scores in total. We consider these $8,001$ alignment scores as the population and refer to the alignments scores between epigenomes of the same cell type (i.e., ESC) as the Group A, and the whole population as the Group B. As there are 8 ESC epigenomes, we obtain 28 alignment scores in the Group A and then calculate the percentile of each of these 28 scores in the population. As shown in Figure S3, 11 out of the 28 scores are among the upper 5% percentile, and the average percentile of alignment scores in group A is 0.256. We also perform a one-tailed t-test to compare the alignment scores of the two groups. The $p$-value of the test is 0.001, which suggests that the chromatin states corresponding to the ESC-associated gene *ANAPC1* have more similar patterns among ESC samples in comparison with the other cell types in the Roadmap dataset. We also use the alignment scores based on the naïve method and repeat all the analysis above.

From Figure S3, we can see that compared to the naïve method, EpiAlign can better distinguish alignment scores among ESCs from others. Similar results are observed for brain and heart too. These results indicate that for a specific cell type, EpiAlign is able to detect the similarity of chromatin states of its tissue-associated genes, suggesting that EpiAlign may be used to differentiate a given cell type from the other tissue and cell types, by evaluating the similarity of epigenetic signals on its associated genes. Also, these results show that EpiAlign can be used to identify tissue-associated regions.

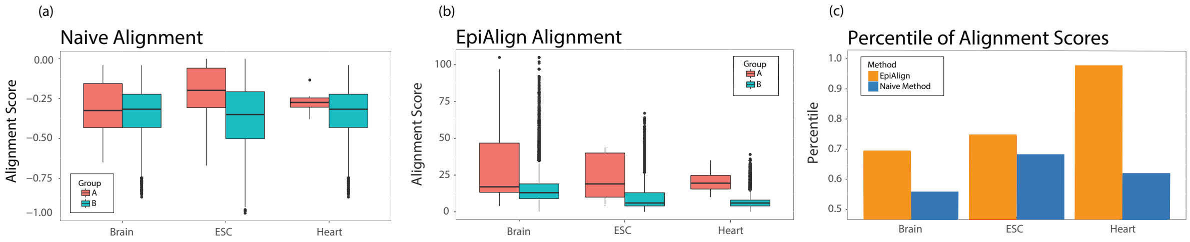


Figure S3: (a)-(b) Boxplots of pairwise alignment scores of chromatin state sequences of a tissue-associated gene within same the cell type (Group A) and across all samples (Group B). We choose an ESC-associated gene *ANAPC1*, a brain-associated gene *AK5*, and a heart-associated gene *ACTN2*. (a) shows the alignment scores by the naïve method, and (b) shows the EpiAlign alignment scores. (c) shows the average percentile of group A scores in Group B for each alignment method.

We also perform hierarchical clustering of the 127 cells using chromatin state sequences of multiple tissue-associated genes. For example, for each of the 118 ESC-associated genes, we use EpiAlign to calculate all the pairwise alignment scores and form a $127 \times 127$ score matrix. We then normalize the scores by dividing the maximum so that for each gene $i$, we get a $127 \times 127$ normalized comparison matrix $M^i$. Then the final distance matrix $D$ is calculated as $D_{jk} = -\sqrt{\sum_{i=1}^{118} M_{jk}^i}$. Finally, we perform complete-linkage hierarchical clustering on the 127 epigenomes based on the distance matrix. The heatmap of the distance matrix and the clustering results are shown in Figure S4(b). The heatmap can roughly distinguish ESC samples from the other cell types. In hierarchical clustering, 7 out of 8 ESC samples are successfully grouped together when the cluster number is set as 10. Similarly, we perform the above analysis based on brain-associated genes, and the heatmap is shown in S4(a). The brain samples can be clearly differentiated. In addition, all the 10 brain samples are successfully grouped together by hierarchical clustering when cluster number is set as 10. The above results confirm EpiAlign's capability to search for similar chromatin state patterns and suggest that chromatin states of the same genomic region are more similar within cell types.

### 8 Male-vs-female vertical alignment

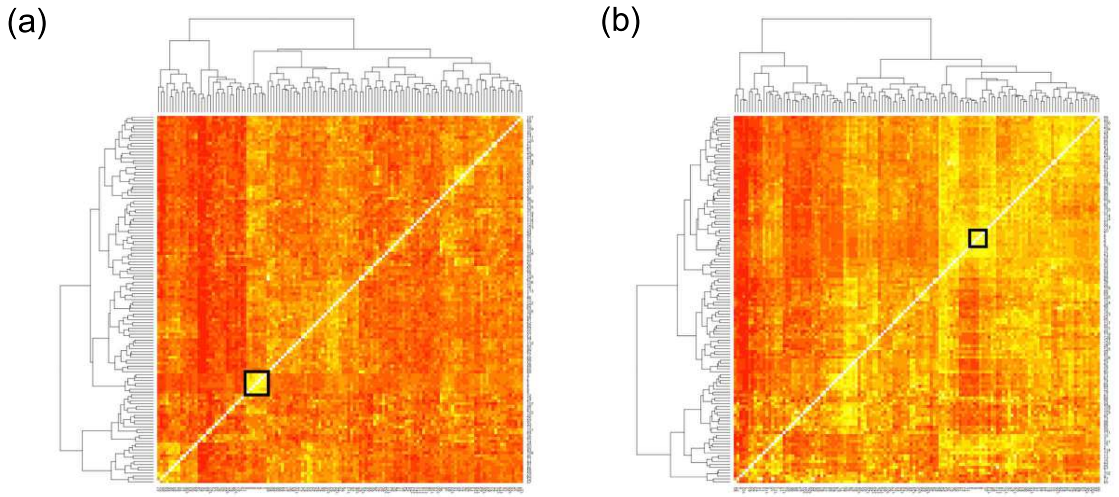


Figure S4: Clustering results using (a) brain-associated genes or (b) ESC-associated genes. Samples in black boxes are (a) brain samples and (b) ESC samples.

### 9 Brain-vs-heart vertical alignment

#### 9.1 Genes with the smallest p-values from one-sided Wilcoxon test

We perform the two-sample one-sided Wilcoxon test between the brain-vs-brain alignment scores and the brain-vs-heart alignment scores for all the protein-coding genes. The top 200 genes that we use to perform the gene ontology enrichment analysis are listed in Table S2.

| | Gene.stable.ID | Chromosome | Gene.name | Strand | Gene.start..bp. | Gene.end..bp. |
|---|---|---|---|---|---|---|
| 1 | ENSG00000004700 | 12 | RECQL | -1 | 21468911 | 21501669 |
| 2 | ENSG00000006047 | 17 | YBX2 | -1 | 7288252 | 7294615 |
| 3 | ENSG00000015592 | 8 | STMN4 | -1 | 27235323 | 27258420 |
| 4 | ENSG00000033122 | 1 | LRRC7 | 1 | 69568398 | 70151945 |
| 5 | ENSG00000034053 | 15 | APBA2 | 1 | 28884483 | 29118315 |
| 6 | ENSG00000047365 | 4 | ARAP2 | -1 | 35948221 | 36244509 |
| 7 | ENSG00000050438 | 12 | SLC4A8 | 1 | 51391317 | 51515763 |
| 8 | ENSG00000054282 | 1 | SDCCAG8 | 1 | 243256034 | 243500092 |
| 9 | ENSG00000056487 | 22 | PHF21B | -1 | 44881162 | 45009999 |
| 10 | ENSG00000064270 | 16 | ATP2C2 | 1 | 84368527 | 84464187 |
| 11 | ENSG00000065609 | 6 | SNAP91 | -1 | 83552880 | 83709691 |
| 12 | ENSG00000066468 | 10 | FGFR2 | -1 | 121478334 | 121598458 |
| 13 | ENSG00000067221 | 15 | STOML1 | -1 | 73978923 | 73994622 |
| 14 | ENSG00000067798 | 12 | NAV3 | 1 | 77324641 | 78213008 |
| 15 | ENSG00000070501 | 8 | POLB | 1 | 42338454 | 42371808 |
| 16 | ENSG00000073803 | 3 | MAP3K13 | 1 | 185282941 | 185489097 |
| 17 | ENSG00000074211 | 4 | PPP2R2C | -1 | 6320578 | 6563600 |
| 18 | ENSG00000077009 | 19 | NMRK2 | 1 | 3933103 | 3942416 |
| 19 | ENSG00000078295 | 5 | ADCY2 | 1 | 7396208 | 7830081 |
| 20 | ENSG00000078725 | 9 | BRINP1 | -1 | 119153458 | 119369467 |
| 21 | ENSG00000084628 | 1 | NKAIN1 | -1 | 31179745 | 31239554 |

| | | | | | | |
|---|---|---|---|---|---|---|
| 22 | ENSG00000088766 | 20 | CRLS1 | 1 | 6006090 | 6040053 |
| 23 | ENSG00000089225 | 12 | TBX5 | -1 | 114353931 | 114408442 |
| 24 | ENSG00000091129 | 7 | NRCAM | -1 | 108147623 | 108456717 |
| 25 | ENSG00000095397 | 9 | WHRN | -1 | 114402080 | 114505450 |
| 26 | ENSG00000100290 | 22 | BIK | 1 | 43110748 | 43129712 |
| 27 | ENSG00000100433 | 14 | KCNK10 | -1 | 88180103 | 88326907 |
| 28 | ENSG00000100505 | 14 | TRIM9 | -1 | 50975262 | 51096061 |
| 29 | ENSG00000102383 | X | ZDHHC15 | -1 | 75368427 | 75523502 |
| 30 | ENSG00000104112 | 15 | SCG3 | 1 | 51681353 | 51721031 |
| 31 | ENSG00000104833 | 19 | TUBB4A | -1 | 6494319 | 6502848 |
| 32 | ENSG00000105048 | 19 | TNNT1 | -1 | 55132794 | 55149354 |
| 33 | ENSG00000106780 | 9 | MEGF9 | -1 | 120600813 | 120714470 |
| 34 | ENSG00000107438 | 10 | PDLIM1 | -1 | 95237572 | 95291024 |
| 35 | ENSG00000108001 | 10 | EBF3 | -1 | 129835283 | 129963841 |
| 36 | ENSG00000108187 | 10 | PBLD | -1 | 68282660 | 68333049 |
| 37 | ENSG00000108688 | 17 | CCL7 | 1 | 34270221 | 34272242 |
| 38 | ENSG00000108830 | 17 | RND2 | 1 | 43025241 | 43032036 |
| 39 | ENSG00000109472 | 4 | CPE | 1 | 165361194 | 165498320 |
| 40 | ENSG00000109654 | 4 | TRIM2 | 1 | 153152342 | 153339320 |
| 41 | ENSG00000109956 | 11 | B3GAT1 | -1 | 134378504 | 134411918 |
| 42 | ENSG00000110042 | 11 | DTX4 | 1 | 59171430 | 59208587 |
| 43 | ENSG00000110076 | 11 | NRXN2 | -1 | 64606174 | 64723188 |
| 44 | ENSG00000110628 | 11 | SLC22A18 | 1 | 2899721 | 2925246 |
| 45 | ENSG00000111605 | 12 | CPSF6 | 1 | 69239537 | 69274358 |
| 46 | ENSG00000111726 | 12 | CMAS | 1 | 22046174 | 22065674 |
| 47 | ENSG00000112041 | 6 | TULP1 | -1 | 35497874 | 35512938 |
| 48 | ENSG00000112139 | 6 | MDGA1 | -1 | 37630679 | 37699306 |
| 49 | ENSG00000112290 | 6 | WASF1 | -1 | 110099819 | 110180004 |
| 50 | ENSG00000112379 | 6 | ARFGEF3 | 1 | 138161921 | 138344663 |
| 51 | ENSG00000113456 | 5 | RAD1 | -1 | 34905264 | 34918989 |
| 52 | ENSG00000113460 | 5 | BRIX1 | 1 | 34915376 | 34925996 |
| 53 | ENSG00000113645 | 5 | WWC1 | 1 | 168291651 | 168472303 |
| 54 | ENSG00000115041 | 2 | KCNIP3 | 1 | 95297304 | 95386083 |
| 55 | ENSG00000115239 | 2 | ASB3 | -1 | 53532672 | 53787610 |
| 56 | ENSG00000117020 | 1 | AKT3 | -1 | 243488233 | 243851079 |
| 57 | ENSG00000117595 | 1 | IRF6 | -1 | 209785623 | 209806175 |
| 58 | ENSG00000118322 | 5 | ATP10B | -1 | 160563120 | 160852214 |
| 59 | ENSG00000120937 | 1 | NPPB | -1 | 11857464 | 11858931 |
| 60 | ENSG00000120963 | 8 | ZNF706 | -1 | 101177878 | 101206193 |
| 61 | ENSG00000121058 | 17 | COIL | -1 | 56938187 | 56961054 |
| 62 | ENSG00000121743 | 13 | GJA3 | -1 | 20138255 | 20161049 |
| 63 | ENSG00000121904 | 1 | CSMD2 | -1 | 33513999 | 34165842 |
| 64 | ENSG00000123560 | X | PLP1 | 1 | 103773718 | 103792619 |
| 65 | ENSG00000124641 | 6 | MED20 | -1 | 41905354 | 41921139 |
| 66 | ENSG00000127955 | 7 | GNAI1 | 1 | 79768028 | 80226181 |
| 67 | ENSG00000128524 | 7 | ATP6V1F | 1 | 128862826 | 128865844 |
| 68 | ENSG00000129250 | 17 | KIF1C | 1 | 4997948 | 5028401 |
| 69 | ENSG00000129991 | 19 | TNNI3 | -1 | 55151767 | 55157773 |

| | | | | | | |
|---|---|---|---|---|---|---|
| 70 | ENSG00000130176 | 19 | CNN1 | 1 | 11538717 | 11550323 |
| 71 | ENSG00000130226 | 7 | DPP6 | 1 | 153887097 | 154894285 |
| 72 | ENSG00000130475 | 19 | FCHO1 | 1 | 17747718 | 17788568 |
| 73 | ENSG00000131409 | 19 | LRRC4B | -1 | 50516892 | 50568045 |
| 74 | ENSG00000131437 | 5 | KIF3A | -1 | 132692628 | 132737638 |
| 75 | ENSG00000132549 | 8 | VPS13B | 1 | 99013266 | 99877580 |
| 76 | ENSG00000133216 | 1 | EPHB2 | 1 | 22710839 | 22921500 |
| 77 | ENSG00000133958 | 14 | UNC79 | 1 | 93333219 | 93707876 |
| 78 | ENSG00000135069 | 9 | PSAT1 | 1 | 78297143 | 78330093 |
| 79 | ENSG00000135269 | 7 | TES | 1 | 116210493 | 116258783 |
| 80 | ENSG00000135298 | 6 | ADGRB3 | 1 | 68635367 | 69389511 |
| 81 | ENSG00000136155 | 13 | SCEL | 1 | 77535674 | 77645263 |
| 82 | ENSG00000136193 | 7 | SCRN1 | -1 | 29920103 | 29990289 |
| 83 | ENSG00000136574 | 8 | GATA4 | 1 | 11676959 | 11760002 |
| 84 | ENSG00000137266 | 6 | SLC22A23 | -1 | 3268962 | 3457022 |
| 85 | ENSG00000139364 | 12 | TMEM132B | 1 | 125186836 | 125662377 |
| 86 | ENSG00000140937 | 16 | CDH11 | -1 | 64943753 | 65126112 |
| 87 | ENSG00000141448 | 18 | GATA6 | 1 | 22169443 | 22202528 |
| 88 | ENSG00000141574 | 17 | SECTM1 | -1 | 82321024 | 82334074 |
| 89 | ENSG00000141738 | 17 | GRB7 | 1 | 39737927 | 39747291 |
| 90 | ENSG00000142949 | 1 | PTPRF | 1 | 43525187 | 43623666 |
| 91 | ENSG00000143951 | 2 | WDPCP | -1 | 63121383 | 63827843 |
| 92 | ENSG00000144369 | 2 | FAM171B | 1 | 186693971 | 186765965 |
| 93 | ENSG00000144857 | 3 | BOC | 1 | 113211003 | 113287459 |
| 94 | ENSG00000145284 | 4 | SCD5 | -1 | 82629539 | 82798857 |
| 95 | ENSG00000145555 | 5 | MYO10 | -1 | 16661914 | 16936276 |
| 96 | ENSG00000145794 | 5 | MEGF10 | 1 | 127290831 | 127465737 |
| 97 | ENSG00000146005 | 5 | PSD2 | 1 | 139795821 | 139844466 |
| 98 | ENSG00000146352 | 6 | CLVS2 | 1 | 122995971 | 123072927 |
| 99 | ENSG00000147488 | 8 | ST18 | -1 | 52110839 | 52460959 |
| 100 | ENSG00000147724 | 8 | FAM135B | -1 | 138130023 | 138496822 |
| 101 | ENSG00000147799 | 8 | ARHGAP39 | -1 | 144529179 | 144605816 |
| 102 | ENSG00000148123 | 9 | PLPPR1 | 1 | 101028709 | 101325135 |
| 103 | ENSG00000149571 | 11 | KIRREL3 | -1 | 126423359 | 127003460 |
| 104 | ENSG00000149596 | 20 | JPH2 | -1 | 44111695 | 44187578 |
| 105 | ENSG00000150477 | 18 | KIAA1328 | 1 | 36829106 | 37232172 |
| 106 | ENSG00000150625 | 4 | GPM6A | -1 | 175632934 | 176002664 |
| 107 | ENSG00000152578 | 11 | GRIA4 | 1 | 105609994 | 105982092 |
| 108 | ENSG00000154229 | 17 | PRKCA | 1 | 66302636 | 66810743 |
| 109 | ENSG00000155886 | 9 | SLC24A2 | -1 | 19507452 | 19786928 |
| 110 | ENSG00000156475 | 5 | PPP2R2B | -1 | 146581146 | 147084784 |
| 111 | ENSG00000157103 | 3 | SLC6A1 | 1 | 10992186 | 11039249 |
| 112 | ENSG00000157423 | 16 | HYDIN | -1 | 70807378 | 71230722 |
| 113 | ENSG00000157851 | 2 | DPYSL5 | 1 | 26847747 | 26950351 |
| 114 | ENSG00000158014 | 1 | SLC30A2 | -1 | 26037252 | 26046133 |
| 115 | ENSG00000158615 | 1 | PPP1R15B | -1 | 204403387 | 204411791 |
| 116 | ENSG00000162706 | 1 | CADM3 | 1 | 159171609 | 159203313 |
| 117 | ENSG00000163449 | 2 | TMEM169 | 1 | 216081866 | 216102783 |

| | | | | | | |
|---|---|---|---|---|---|---|
| 118 | ENSG00000164107 | 4 | HAND2 | -1 | 173524969 | 173530229 |
| 119 | ENSG00000164163 | 4 | ABCE1 | 1 | 145097932 | 145129179 |
| 120 | ENSG00000164532 | 7 | TBX20 | -1 | 35202430 | 35254147 |
| 121 | ENSG00000164542 | 7 | KIAA0895 | -1 | 36324221 | 36390125 |
| 122 | ENSG00000165312 | 10 | OTUD1 | 1 | 23439458 | 23442390 |
| 123 | ENSG00000165527 | 14 | ARF6 | 1 | 49893092 | 49897054 |
| 124 | ENSG00000165548 | 14 | TMEM63C | 1 | 77116568 | 77259495 |
| 125 | ENSG00000165566 | 13 | AMER2 | -1 | 25161684 | 25172288 |
| 126 | ENSG00000166501 | 16 | PRKCB | 1 | 23835946 | 24220611 |
| 127 | ENSG00000166831 | 15 | RBPMS2 | -1 | 64739892 | 64775587 |
| 128 | ENSG00000166922 | 15 | SCG5 | 1 | 32641676 | 32697098 |
| 129 | ENSG00000167553 | 12 | TUBA1C | 1 | 49188736 | 49274603 |
| 130 | ENSG00000168280 | 2 | KIF5C | 1 | 148875250 | 149026759 |
| 131 | ENSG00000168495 | 8 | POLR3D | 1 | 22245104 | 22254600 |
| 132 | ENSG00000168958 | 2 | MFF | 1 | 227325151 | 227357836 |
| 133 | ENSG00000170091 | 5 | NSG2 | 1 | 174045604 | 174243501 |
| 134 | ENSG00000170185 | 4 | USP38 | 1 | 143184917 | 143223830 |
| 135 | ENSG00000171954 | 19 | CYP4F22 | 1 | 15508493 | 15552317 |
| 136 | ENSG00000172379 | 15 | ARNT2 | 1 | 80404350 | 80597937 |
| 137 | ENSG00000172461 | 6 | FUT9 | 1 | 96015984 | 96215612 |
| 138 | ENSG00000172995 | 3 | ARPP21 | 1 | 35638945 | 35794496 |
| 139 | ENSG00000173530 | 8 | TNFRSF10D | -1 | 23135588 | 23164030 |
| 140 | ENSG00000173898 | 11 | SPTBN2 | -1 | 66685248 | 66729226 |
| 141 | ENSG00000174099 | 12 | MSRB3 | 1 | 65278643 | 65491430 |
| 142 | ENSG00000174407 | 20 | MIR1-1HG | 1 | 62550453 | 62570764 |
| 143 | ENSG00000174672 | 11 | BRSK2 | 1 | 1389899 | 1462689 |
| 144 | ENSG00000175084 | 2 | DES | 1 | 219418377 | 219426739 |
| 145 | ENSG00000175087 | 1 | PDIK1L | 1 | 26111165 | 26125543 |
| 146 | ENSG00000175161 | 3 | CADM2 | 1 | 84958981 | 86074429 |
| 147 | ENSG00000176049 | 5 | JAKMIP2 | -1 | 147585439 | 147782848 |
| 148 | ENSG00000177103 | 11 | DSCAML1 | -1 | 117427773 | 117817525 |
| 149 | ENSG00000177508 | 16 | IRX3 | -1 | 54283304 | 54286763 |
| 150 | ENSG00000177807 | 1 | KCNJ10 | -1 | 159998651 | 160070483 |
| 151 | ENSG00000178445 | 9 | GLDC | -1 | 6532464 | 6645783 |
| 152 | ENSG00000179242 | 20 | CDH4 | 1 | 61252426 | 61940617 |
| 153 | ENSG00000179314 | 17 | WSCD1 | 1 | 6057807 | 6124427 |
| 154 | ENSG00000179915 | 2 | NRXN1 | -1 | 49918505 | 51225575 |
| 155 | ENSG00000180287 | 1 | PLD5 | -1 | 242082986 | 242524696 |
| 156 | ENSG00000182600 | 2 | SNORC | 1 | 232857270 | 232878708 |
| 157 | ENSG00000183072 | 5 | NKX2-5 | -1 | 173232109 | 173235357 |
| 158 | ENSG00000185155 | 1 | MIXL1 | 1 | 226223618 | 226227054 |
| 159 | ENSG00000185156 | 17 | MFSD6L | -1 | 8797162 | 8799349 |
| 160 | ENSG00000185565 | 3 | LSAMP | -1 | 115802363 | 117139389 |
| 161 | ENSG00000185627 | 11 | PSMD13 | 1 | 236546 | 252984 |
| 162 | ENSG00000185818 | 4 | NAT8L | 1 | 2059512 | 2069089 |
| 163 | ENSG00000185973 | X | TMLHE | -1 | 155490115 | 155669944 |
| 164 | ENSG00000186231 | 6 | KLHL32 | 1 | 96924620 | 97140754 |
| 165 | ENSG00000187164 | 10 | SHTN1 | -1 | 116881482 | 117126586 |

| | | | | | | |
|---|---|---|---|---|---|---|
| 166 | ENSG00000187634 | 1 | SAMD11 | 1 | 923928 | 944581 |
| 167 | ENSG00000188015 | 1 | S100A3 | -1 | 153547329 | 153549372 |
| 168 | ENSG00000188316 | 10 | ENO4 | 1 | 116849512 | 116911788 |
| 169 | ENSG00000188522 | 17 | FAM83G | -1 | 18968789 | 19004804 |
| 170 | ENSG00000196220 | 3 | SRGAP3 | -1 | 8980591 | 9363053 |
| 171 | ENSG00000196338 | X | NLGN3 | 1 | 71144831 | 71171201 |
| 172 | ENSG00000196361 | 19 | ELAVL3 | -1 | 11451326 | 11481046 |
| 173 | ENSG00000196376 | 6 | SLC35F1 | 1 | 117907526 | 118317676 |
| 174 | ENSG00000196581 | 1 | AJAP1 | 1 | 4654732 | 4792534 |
| 175 | ENSG00000196628 | 18 | TCF4 | -1 | 55222331 | 55664787 |
| 176 | ENSG00000196767 | X | POU3F4 | 1 | 83508261 | 83512127 |
| 177 | ENSG00000197728 | 12 | RPS26 | 1 | 56041351 | 56044697 |
| 178 | ENSG00000198216 | 1 | CACNA1E | 1 | 181317690 | 181808084 |
| 179 | ENSG00000198513 | 14 | ATL1 | 1 | 50532509 | 50633068 |
| 180 | ENSG00000198732 | 14 | SMOC1 | 1 | 69854131 | 70032366 |
| 181 | ENSG00000203930 | X | LINC00632 | 1 | 140709562 | 140793215 |
| 182 | ENSG00000204011 | 9 | COL5A1-AS1 | -1 | 134649385 | 134652843 |
| 183 | ENSG00000204344 | 6 | STK19 | 1 | 31971091 | 31982821 |
| 184 | ENSG00000204624 | 1 | DISP3 | 1 | 11479166 | 11537584 |
| 185 | ENSG00000204683 | 10 | C10orf113 | -1 | 21125763 | 21146559 |
| 186 | ENSG00000205758 | 21 | CRYZL1 | -1 | 33589341 | 33643926 |
| 187 | ENSG00000213578 | 15 | CPLX3 | 1 | 74826547 | 74831802 |
| 188 | ENSG00000214160 | 3 | ALG3 | -1 | 184242301 | 184249548 |
| 189 | ENSG00000214338 | 6 | SOGA3 | -1 | 127472794 | 127519191 |
| 190 | ENSG00000214595 | 2 | EML6 | 1 | 54723499 | 54972025 |
| 191 | ENSG00000219438 | 22 | FAM19A5 | 1 | 48489460 | 48850912 |
| 192 | ENSG00000221818 | 8 | EBF2 | -1 | 25841730 | 26045397 |
| 193 | ENSG00000235568 | 22 | NFAM1 | -1 | 42380410 | 42432395 |
| 194 | ENSG00000237330 | 1 | RNF223 | -1 | 1070966 | 1074307 |
| 195 | ENSG00000241370 | 6 | RPP21 | 1 | 30345131 | 30346884 |
| 196 | ENSG00000243232 | 5 | PCDHAC2 | 1 | 140966235 | 141012344 |
| 197 | ENSG00000243449 | 4 | C4orf48 | 1 | 2041993 | 2043970 |
| 198 | ENSG00000248383 | 5 | PCDHAC1 | 1 | 140926369 | 141012344 |
| 199 | ENSG00000253276 | 7 | CCDC71L | -1 | 106656765 | 106660996 |
| 200 | ENSG00000255537 | 11 | AP000708.1 | 1 | 125495214 | 125499528 |

Table S2: Top 200 significant genes from the one-sided Wilcoxon test that compares brain-vs-brain scores and brain-vs-heart scores.

#### 9.2 Tissue-specific genes receive lower $p$-values in Wilcoxon test

From the boxplots in Figure S5, we can see that brain-specific genes and heart-specific genes receive lower $p$-values from the one-sided Wilcoxon test that compares brain-vs-brain alignment scores and brain-vs-heart alignment scores. When comparing heart-vs-heart scores and heart-vs-brain scores, heart-specific genes have much lower $p$-values. These results indicates that EpiAlign can correctly capture cell-type-characteristic chromatin state patterns.

| | Gene.stable.ID | Gene.name | Chromosome | Gene.description |
|---|---|---|---|---|
| 1 | ENSG00000012660 | ELOVL5 | 6 | ELOVL fatty acid elongase 5 |
| 2 | ENSG00000111832 | RWDD1 | 6 | RWD domain containing 1 |
| 3 | ENSG00000112232 | KHDRBS2 | 6 | KH RNA binding domain containing, signal transduction associated 2 |
| 4 | ENSG00000135968 | GCC2 | 2 | GRIP and coiled-coil domain containing 2 |
| 5 | ENSG00000136485 | DCAF7 | 17 | DDB1 and CUL4 associated factor 7 |
| 6 | ENSG00000138035 | PNPT1 | 2 | polyribonucleotide nucleotidyltransferase 1 |
| 7 | ENSG00000138398 | PPIG | 2 | peptidylprolyl isomerase G |
| 8 | ENSG00000139053 | PDE6H | 12 | phosphodiesterase 6H |
| 9 | ENSG00000168958 | MFF | 2 | mitochondrial fission factor |
| 10 | ENSG00000170293 | CMTM8 | 3 | CKLF like MARVEL transmembrane domain containing 2 |
| 11 | ENSG00000173572 | NLRP13 | 19 | NLR family pyrin domain containing 13 |
| 12 | ENSG00000197360 | ZNF98 | 19 | zinc finger protein 98 |

Table S1: Genes not on chromosome X among the top 200 genes with the smallest $p$-values from comparing male-vs-male scores and male-vs-female scores.

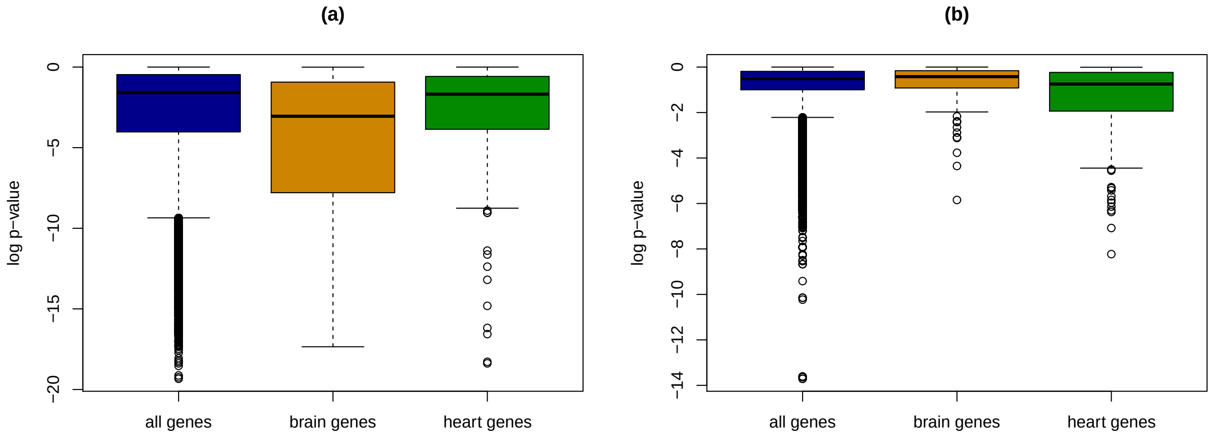


Figure S5: Boxplots of $p$-values of all genes, brain-specific genes and heart-specific genes from Wilcoxon tests which compares (a) brain-vs-brain scores with brain-vs-heart scores or (b) heart-vs-heart scores with brain-vs-heart scores.

### 10 Motif analysis

The top 200 regions with the highest horizontal alignment scores and the cluster index to which each region belongs are listed in Table S3. We use the motif-discovery tool MEME and find that all the four clusters are characterized by certain motifs, the top motifs reported by MEME are:
Cluster 1: “ihihihihihihihihihio”; “edegbabg”; “aehihedhdeo”.
Cluster 2: “gegegogegegogogegege”; “gegegegegegegegegege”; gegegegegegegegegege”;
Cluster 3: “goglkjklnog”; “dededededededededede”; “gegegegedegededegege”
Cluster 4: “oaogegegegbgege”; “mlklmlmlklklklk”; “nogogogo”.

| | sample | chromosome | start.position | end.position | cluster |
|---|---|---|---|---|---|
| 1 | E003 | chr22 | 22800001 | 23300000 | 1 |
| 2 | E003 | chr21 | 10700001 | 11200000 | 1 |
| 3 | E003 | chrX | 61500001 | 62000000 | 1 |
| 4 | E003 | chr17 | 46400001 | 46900000 | 1 |
| 5 | E003 | chr20 | 59400001 | 59900000 | 1 |
| 6 | E003 | chr12 | 54100001 | 54600000 | 1 |
| 7 | E003 | chr12 | 37900001 | 38400000 | 1 |
| 8 | E003 | chr7 | 26900001 | 27400000 | 1 |
| 9 | E003 | chr2 | 176700001 | 177200000 | 1 |
| 10 | E003 | chr3 | 61500001 | 62000000 | 1 |
| 11 | E003 | chr6 | 157100001 | 157600000 | 1 |
| 12 | E003 | chr8 | 43400001 | 43900000 | 1 |
| 13 | E003 | chr16 | 33800001 | 34300000 | 1 |
| 14 | E003 | chr15 | 99100001 | 99600000 | 1 |
| 15 | E003 | chr11 | 2300001 | 2800000 | 1 |
| 16 | E003 | chrY | 9900001 | 10400000 | 1 |
| 17 | E003 | chr2 | 92200001 | 92700000 | 1 |
| 18 | E003 | chr8 | 128600001 | 129100000 | 1 |
| 19 | E003 | chr7 | 61700001 | 62200000 | 2 |
| 20 | E003 | chr7 | 101300001 | 101800000 | 2 |
| 21 | E003 | chr5 | 89900001 | 90400000 | 2 |
| 22 | E003 | chr22 | 22300001 | 22800000 | 2 |
| 23 | E003 | chr4 | 78900001 | 79400000 | 2 |
| 24 | E003 | chr18 | 60100001 | 60600000 | 2 |
| 25 | E003 | chr10 | 34600001 | 35100000 | 2 |
| 26 | E003 | chr16 | 49400001 | 49900000 | 2 |
| 27 | E003 | chr6 | 15200001 | 15700000 | 2 |
| 28 | E003 | chr22 | 33900001 | 34400000 | 2 |
| 29 | E003 | chr5 | 54300001 | 54800000 | 2 |
| 30 | E003 | chr10 | 42200001 | 42700000 | 2 |
| 31 | E003 | chr14 | 89700001 | 90200000 | 2 |
| 32 | E003 | chr2 | 153200001 | 153700000 | 2 |
| 33 | E003 | chr2 | 121300001 | 121800000 | 2 |
| 34 | E003 | chr1 | 164300001 | 164800000 | 2 |
| 35 | E003 | chr19 | 37500001 | 38000000 | 2 |
| 36 | E003 | chr1 | 8400001 | 8900000 | 3 |
| 37 | E003 | chr4 | 190400001 | 190900000 | 2 |
| 38 | E003 | chr1 | 64200001 | 64700000 | 2 |
| 39 | E003 | chr5 | 106600001 | 107100000 | 2 |
| 40 | E003 | chr10 | 12000001 | 12500000 | 2 |
| 41 | E003 | chr5 | 13900001 | 14400000 | 2 |
| 42 | E003 | chr7 | 154900001 | 155400000 | 2 |
| 43 | E003 | chr11 | 31700001 | 32200000 | 2 |
| 44 | E003 | chr4 | 183900001 | 184400000 | 2 |
| 45 | E003 | chr12 | 132600001 | 133100000 | 2 |
| 46 | E003 | chr2 | 55200001 | 55700000 | 2 |

| | | | | | |
|---|---|---|---|---|---|
| 47 | E003 | chr8 | 131200001 | 131700000 | 2 |
| 48 | E003 | chr8 | 142200001 | 142700000 | 2 |
| 49 | E003 | chr17 | 59700001 | 60200000 | 2 |
| 50 | E003 | chr12 | 34400001 | 34900000 | 2 |
| 51 | E003 | chr11 | 12500001 | 13000000 | 2 |
| 52 | E003 | chr15 | 57300001 | 57800000 | 2 |
| 53 | E003 | chr20 | 49200001 | 49700000 | 2 |
| 54 | E003 | chr10 | 114400001 | 114900000 | 2 |
| 55 | E003 | chr5 | 87600001 | 88100000 | 2 |
| 56 | E003 | chr17 | 28800001 | 29300000 | 2 |
| 57 | E003 | chr22 | 29000001 | 29500000 | 2 |
| 58 | E003 | chr7 | 105300001 | 105800000 | 2 |
| 59 | E003 | chr22 | 17600001 | 18100000 | 2 |
| 60 | E003 | chr19 | 12800001 | 13300000 | 2 |
| 61 | E003 | chr12 | 32100001 | 32600000 | 2 |
| 62 | E003 | chr2 | 236200001 | 236700000 | 2 |
| 63 | E003 | chr3 | 185400001 | 185900000 | 2 |
| 64 | E003 | chr12 | 130400001 | 130900000 | 2 |
| 65 | E003 | chr15 | 26800001 | 27300000 | 2 |
| 66 | E003 | chr16 | 46000001 | 46500000 | 2 |
| 67 | E003 | chr11 | 107800001 | 108300000 | 2 |
| 68 | E003 | chr6 | 148900001 | 149400000 | 2 |
| 69 | E003 | chr4 | 93300001 | 93800000 | 2 |
| 70 | E003 | chr9 | 128200001 | 128700000 | 2 |
| 71 | E003 | chr15 | 50600001 | 51100000 | 2 |
| 72 | E003 | chr17 | 3900001 | 4400000 | 2 |
| 73 | E003 | chr1 | 235200001 | 235700000 | 2 |
| 74 | E003 | chr9 | 140300001 | 140800000 | 2 |
| 75 | E003 | chr19 | 1400001 | 1900000 | 2 |
| 76 | E003 | chr10 | 88400001 | 88900000 | 2 |
| 77 | E003 | chr15 | 28200001 | 28700000 | 2 |
| 78 | E003 | chr3 | 31600001 | 32100000 | 2 |
| 79 | E003 | chr2 | 188900001 | 189400000 | 2 |
| 80 | E003 | chr22 | 31800001 | 32300000 | 2 |
| 81 | E003 | chr15 | 44600001 | 45100000 | 2 |
| 82 | E003 | chr4 | 85400001 | 85900000 | 2 |
| 83 | E003 | chr13 | 98700001 | 99200000 | 2 |
| 84 | E003 | chr13 | 28300001 | 28800000 | 2 |
| 85 | E003 | chr9 | 33100001 | 33600000 | 2 |
| 86 | E003 | chr15 | 63800001 | 64300000 | 2 |
| 87 | E003 | chr1 | 39500001 | 40000000 | 2 |
| 88 | E003 | chr10 | 80500001 | 81000000 | 2 |
| 89 | E003 | chr3 | 121100001 | 121600000 | 2 |
| 90 | E003 | chr13 | 41000001 | 41500000 | 2 |
| 91 | E003 | chr16 | 89400001 | 89900000 | 2 |
| 92 | E003 | chr2 | 32400001 | 32900000 | 2 |
| 93 | E003 | chr1 | 219900001 | 220400000 | 2 |
| 94 | E003 | chr2 | 119400001 | 119900000 | 2 |

| | | | | | |
|---|---|---|---|---|---|
| 95 | E003 | chr1 | 10500001 | 11000000 | 2 |
| 96 | E003 | chr1 | 200200001 | 200700000 | 2 |
| 97 | E003 | chr12 | 112300001 | 112800000 | 2 |
| 98 | E003 | chr14 | 99600001 | 100100000 | 2 |
| 99 | E003 | chr8 | 30000001 | 30500000 | 2 |
| 100 | E003 | chr10 | 74800001 | 75300000 | 2 |
| 101 | E003 | chr13 | 113300001 | 113800000 | 2 |
| 102 | E003 | chr10 | 102800001 | 103300000 | 2 |
| 103 | E003 | chr18 | 52900001 | 53400000 | 2 |
| 104 | E003 | chr15 | 59300001 | 59800000 | 2 |
| 105 | E003 | chr16 | 1400001 | 1900000 | 2 |
| 106 | E003 | chr16 | 81200001 | 81700000 | 2 |
| 107 | E003 | chr8 | 102500001 | 103000000 | 2 |
| 108 | E003 | chr6 | 56300001 | 56800000 | 2 |
| 109 | E003 | chr13 | 100200001 | 100700000 | 2 |
| 110 | E003 | chr2 | 109000001 | 109500000 | 2 |
| 111 | E003 | chr10 | 126400001 | 126900000 | 2 |
| 112 | E003 | chr8 | 46800001 | 47300000 | 2 |
| 113 | E003 | chr11 | 126200001 | 126700000 | 2 |
| 114 | E003 | chr6 | 41100001 | 41600000 | 3 |
| 115 | E003 | chr2 | 102300001 | 102800000 | 3 |
| 116 | E003 | chr1 | 233000001 | 233500000 | 3 |
| 117 | E003 | chr9 | 16400001 | 16900000 | 3 |
| 118 | E003 | chr21 | 40200001 | 40700000 | 3 |
| 119 | E003 | chr9 | 130800001 | 131300000 | 3 |
| 120 | E003 | chr17 | 2600001 | 3100000 | 3 |
| 121 | E003 | chr10 | 96800001 | 97300000 | 3 |
| 122 | E003 | chrX | 16700001 | 17200000 | 3 |
| 123 | E003 | chr13 | 100700001 | 101200000 | 3 |
| 124 | E003 | chr3 | 65500001 | 66000000 | 3 |
| 125 | E003 | chr17 | 25400001 | 25900000 | 3 |
| 126 | E003 | chr15 | 40100001 | 40600000 | 3 |
| 127 | E003 | chr5 | 64800001 | 65300000 | 3 |
| 128 | E003 | chr9 | 94800001 | 95300000 | 3 |
| 129 | E003 | chr9 | 124000001 | 124500000 | 3 |
| 130 | E003 | chr17 | 55300001 | 55800000 | 3 |
| 131 | E003 | chr22 | 43300001 | 43800000 | 3 |
| 132 | E003 | chr1 | 155500001 | 156000000 | 2 |
| 133 | E003 | chr9 | 125400001 | 125900000 | 2 |
| 134 | E003 | chr4 | 184500001 | 185000000 | 2 |
| 135 | E003 | chr20 | 35600001 | 36100000 | 3 |
| 136 | E003 | chr1 | 17700001 | 18200000 | 3 |
| 137 | E003 | chr9 | 70100001 | 70600000 | 3 |
| 138 | E003 | chr8 | 97400001 | 97900000 | 3 |
| 139 | E003 | chr22 | 40400001 | 40900000 | 3 |
| 140 | E003 | chr8 | 141600001 | 142100000 | 3 |
| 141 | E003 | chr12 | 11600001 | 12100000 | 3 |
| 142 | E003 | chr9 | 37600001 | 38100000 | 3 |

| | | | | | |
|---|---|---|---|---|---|
| 143 | E003 | chr8 | 102000001 | 102500000 | 3 |
| 144 | E003 | chr9 | 23700001 | 24200000 | 3 |
| 145 | E003 | chr22 | 45200001 | 45700000 | 3 |
| 146 | E003 | chr3 | 171700001 | 172200000 | 3 |
| 147 | E003 | chr15 | 90900001 | 91400000 | 3 |
| 148 | E003 | chr1 | 161800001 | 162300000 | 3 |
| 149 | E003 | chr15 | 42300001 | 42800000 | 3 |
| 150 | E003 | chr11 | 63600001 | 64100000 | 3 |
| 151 | E003 | chr1 | 21500001 | 22000000 | 3 |
| 152 | E003 | chr1 | 179800001 | 180300000 | 3 |
| 153 | E003 | chr5 | 46000001 | 46500000 | 3 |
| 154 | E003 | chr10 | 79200001 | 79700000 | 3 |
| 155 | E003 | chr18 | 55600001 | 56100000 | 3 |
| 156 | E003 | chr4 | 68300001 | 68800000 | 3 |
| 157 | E003 | chr7 | 121900001 | 122400000 | 3 |
| 158 | E003 | chr17 | 30200001 | 30700000 | 3 |
| 159 | E003 | chr11 | 61300001 | 61800000 | 3 |
| 160 | E003 | chr5 | 70500001 | 71000000 | 3 |
| 161 | E003 | chr2 | 202700001 | 203200000 | 3 |
| 162 | E003 | chr6 | 136500001 | 137000000 | 3 |
| 163 | E003 | chr1 | 23700001 | 24200000 | 3 |
| 164 | E003 | chr2 | 106100001 | 106600000 | 3 |
| 165 | E003 | chr4 | 48900001 | 49400000 | 3 |
| 166 | E003 | chr2 | 183700001 | 184200000 | 3 |
| 167 | E003 | chr6 | 21600001 | 22100000 | 2 |
| 168 | E003 | chr2 | 43400001 | 43900000 | 3 |
| 169 | E003 | chr16 | 72800001 | 73300000 | 3 |
| 170 | E003 | chr19 | 9200001 | 9700000 | 3 |
| 171 | E003 | chr1 | 32100001 | 32600000 | 3 |
| 172 | E003 | chr17 | 15600001 | 16100000 | 3 |
| 173 | E003 | chr17 | 27000001 | 27500000 | 3 |
| 174 | E003 | chr6 | 168200001 | 168700000 | 2 |
| 175 | E003 | chr6 | 37200001 | 37700000 | 2 |
| 176 | E003 | chr11 | 48000001 | 48500000 | 4 |
| 177 | E003 | chr10 | 7900001 | 8400000 | 4 |
| 178 | E003 | chr1 | 12300001 | 12800000 | 4 |
| 179 | E003 | chr9 | 111600001 | 112100000 | 4 |
| 180 | E003 | chr12 | 124900001 | 125400000 | 4 |
| 181 | E003 | chr7 | 55500001 | 56000000 | 4 |
| 182 | E003 | chr16 | 69000001 | 69500000 | 4 |
| 183 | E003 | chr15 | 43000001 | 43500000 | 4 |
| 184 | E003 | chr3 | 47400001 | 47900000 | 4 |
| 185 | E003 | chr10 | 70500001 | 71000000 | 4 |
| 186 | E003 | chr7 | 2400001 | 2900000 | 4 |
| 187 | E003 | chr4 | 7800001 | 8300000 | 4 |
| 188 | E003 | chr11 | 50300001 | 50800000 | 4 |
| 189 | E003 | chr6 | 166800001 | 167300000 | 4 |
| 190 | E003 | chr18 | 74600001 | 75100000 | 4 |

| | | | | | |
|---|---|---|---|---|---|
| 191 | E003 | chr5 | 139800001 | 140300000 | 4 |
| 192 | E003 | chr14 | 77300001 | 77800000 | 4 |
| 193 | E003 | chr11 | 121100001 | 121600000 | 4 |
| 194 | E003 | chr15 | 35000001 | 35500000 | 4 |
| 195 | E003 | chr6 | 29400001 | 29900000 | 4 |
| 196 | E003 | chr5 | 31400001 | 31900000 | 4 |
| 197 | E003 | chr7 | 23000001 | 23500000 | 4 |
| 198 | E003 | chr1 | 47800001 | 48300000 | 4 |
| 199 | E003 | chr10 | 13700001 | 14200000 | 4 |
| 200 | E003 | chr12 | 2900001 | 3400000 | 4 |

Table S3: The top 200 regions with the highest horizontal alignment scores are well partitioned into four clusters by average-linkage hierarchical clustering. The last column of the table is the cluster index to which each region belongs.

#### 11 GO analysis in cross-species application of EpiAlign

We obtain mouse-human homologous gene pairs from Ensembl BioMart and sort the mouse genes with lengths 200-400 kb by gene lengths and divide the homologous gene pairs into 12 groups each with 50 pairs. We look at the molecule function GO terms of the homolog pairs that have the highest alignment scores in each group. The result (Table S4) shows that homolougous genes with high alignment scores are also very similar in molecule function. The result also indicates that EpiAlign can identify homologous genes whose epigenetic patterns are more conserved in evolution, shedding new insights into translating scientific discoveries in mice into humans.

#### 12 Biological discovery based on top 8-mers

[2] We thank the referee for this comment, we have made the following discovery and have revised our manuscript on page 10 to discuss the potential improvement of EpiAlign: We have the following interesting findings when looking at the most common 8-mers in chromatin states identified by ChromHMM: we count the occurrence number of all the 8-mer strings from the epigenome of ESC sample E003. We look at the most frequently occurred 8-mer strings containing active TSS state (represented by 'a'). The most frequent 8-mer is 'aededede', where 'e' represents weak transcript and 'd' represents strong transcript. This 8-mer can be interpreted as active gene region. However, there is also another 8-mer 'oioaoioi', which frequently occurs but is hard to interpret. Here, 'o' represents quiescent/low state and 'i' represents heterochromatin. We select all ESC, heart and brain samples and inspected the overlap between known TSS and these two 8-mers. The results show that in all these samples, a high proportion (71% on average) of the 8-mer 'aededede' discovered have an overlap with known TSS while only a small proportion ( 28% on average) of the 8-mer 'oioaoioi' have an overlap with known TSS. This result indicates that some of the state 'a' in 'oioaoioi' may be noise.

#### References

[1] Yang Yang, Yu-Cheng T Yang, Jiapei Yuan, Zhi John Lu, and Jingyi Jessica Li. Large-scale mapping of mammalian transcriptomes identifies conserved genes associated with different cell

| Homologous pair | | GO: molecule function | |
|---|---|---|---|
| Human | Mouse | Human | Mouse |
| AK5 | AK5 | nucleotide binding | nucleotide binding |
| GABRB2 | GABRB2 | transmembrane signaling receptor activity | transmembrane signaling receptor activity |
| CDH2 | CDH2 | calcium ion binding | calcium ion binding |
| GAB2 | GAB2 | transmembrane receptor protein tyrosine kinase adaptor activity | transmembrane receptor protein tyrosine kinase adaptor activity |
| NEB | NEB | actin binding | actin binding |
| CAMK4 | CAMK4 | nucleotide binding | nucleotide binding |
| PAPPA2 | PAPPA2 | metalloendopeptidase activity | metalloendopeptidase activity |
| CNTN1 | CNTN1 | protein binding | protein binding |
| DNAH7 | DNAH7b | microtubule motor activity | microtubule motor activity |
| TMEM132C | TMEM132C | not available | not available |
| SPATA16 | SPATA16 | not available | not available |
| ADCY2 | ADCY2 | nucleotide binding | nucleotide binding |
| Homologous pair | | GO: biological process | |
| Human | Mouse | Human | Mouse |
| AK5 | AK5 | nucleobase-containing compound metabolic process | nucleobase-containing compound metabolic process |
| GABRB2 | GABRB2 | ion transport | ion transport |
| CDH2 | CDH2 | cell morphogenesis | cell adhesion |
| GAB2 | GAB2 | transmembrane receptor protein tyrosine kinase signaling pathway | transmembrane receptor protein tyrosine kinase signaling pathway |
| NEB | NEB | muscle organ development | regulation of actin filament length |
| CAMK4 | CAMK4 | adaptive immune response | protein phosphorylation |
| PAPPA2 | PAPPA2 | regulation of cell growth | proteolysis |
| CNTN1 | CNTN1 | cell adhesion | cell adhesion |
| DNAH7 | DNAH7b | microtubule-based movement | microtubule-based movement |
| TMEM132C | TMEM132C | not available | not available |
| SPATA16 | SPATA16 | not available | not available |
| ADCY2 | ADCY2 | renal water homeostasis c | AMP biosynthetic process |

Table S4: The top 1 GO terms (both molecule function and biological process) of the homolog pairs that have the highest alignment scores in each group.

states. *Nucleic acids research*, 45(4):1657–1672, 2017.

[2] W James Kent, Charles W Sugnet, Terrence S Furey, Krishna M Roskin, Tom H Pringle, Alan M Zahler, and David Haussler. The human genome browser at ucsc. *Genome research*, 12(6):996–1006, 2002.
